# Elevated ferritin as a companion biomarker of myeloid-driven inflammation in RNP/Sm co-positive, treatment-resistant SLE endotype

**DOI:** 10.64898/2026.02.12.705402

**Authors:** Muhammad RA Shipa, Claire Beesley, Daniel McClusky, Vincent Guichard, Sharon A. Chung, Laura A. Cooney, Derek Gilroy, Michael R Ehrenstein

**Author notes:** **Correspondence to** Dr. Muhammad Shipa, Department of Ageing, Rheumatology and Regenerative Medicine, University College London, London, WC1E 6JF, Prof. Michael Ehrenstein, Department of Ageing, Rheumatology and Regenerative Medicine, University College London, London, WC1E 6JF.

## Abstract

**Background:** The profound molecular heterogeneity of SLE remains a fundamental barrier to therapeutic progress.

**Methods:** We integrated clinical and multi-omic profiling, including transcriptomics, autoantigen microarrays, proteomics, and flow cytometry, across three randomised trials (BEAT-Lupus, CALIBRATE, ACCESS) and two observational cohorts to identify distinct lupus endotypes.

**Findings:** Using multivariate distance-based matching of autoantibody profiles to integrate molecular heterogeneity, we identified anti-RNP/anti-Sm co-positivity (RNP+Sm+) as a distinct myeloid-dominant inflammatory endotype.

Enriched in patients of Black ancestry (approx. 50%), RNP+Sm+ SLE is characterised by expanded intermediate monocytes showing enhanced TLR4-driven inflammation during flares, elevated pro-inflammatory cytokines (TNF-alpha, IL-6, IL-12, IFN-gamma, CCL2), IFNA10-biased interferon signalling, and systemic metabolic activation. IgG autoantibody profiling confirmed epitope spreading to spliceosomes and novel autoreactivity against circadian-metabolic regulators (SIRT1, NCOA1, SREBF1).

Clinically, this endotype manifests as high-grade disease activity (2.5-fold flare risk), nephritis, vasculitis, and enteritis. Hyperferritinaemia correlates with flares exclusively in RNP+Sm+ SLE (r=0.81), reflecting underlying macrophage activation. RNP+Sm+ patients exhibit profound therapeutic resistance: 44% failed first-line immunosuppression (rising to 61% in Black patients), while also showing substantially reduced efficacy (60% lower response) with second-line B-cell-depleting therapies (rituximab, obinutuzumab). Resistance to rituximab therapy was driven by rapid B-cell repopulation, rising BAFF levels, and sustained cytokines despite peripheral depletion. Consequently, they remain heavily steroid-dependent and accrued greater organ damage.

**Conclusion:** The RNP+Sm+ signature defines a high-risk, refractory, myeloid-driven lupus endotype characterised by activity linked hyperferritinemia that likely requires therapies directed at the underlying interferon and myeloid-centred pathways.

**Funding:** BEAT-Lupus: Arthritis UK and GSK. ACCESS and CALIBRATE: National Institute of Allergy and Infectious Diseases of the NIH.

**Context and Significance:** Systemic lupus erythematosus (SLE) is immunologically heterogeneous, yet conventional classifications fail to predict therapeutic outcomes. While autoantibodies are central to diagnosis, whether specific combinations define mechanistically distinct subsets remains unresolved. Here, we demonstrate that anti-RNP/anti-Sm co-positivity (RNP⁺Sm⁺) identifies a clinically aggressive, myeloid-dominant endotype disproportionately affecting Black patients. Multi-omics integration revealed expansion of intermediate monocytes with enhanced TLR4 expression, IFNA10-biased interferon signalling, and metabolic reprogramming. Clinically, RNP⁺Sm⁺ patients exhibit nephritis, vasculitis, higher flare rates, and profound first-line therapeutic resistance. Subsequently, this endotype shows poor response to B-cell-depleting therapies, necessitating prolonged glucocorticoid dependence and accelerated organ damage. Establishing RNP⁺Sm⁺ as a lupus endotype and ferritin as a companion disease activity biomarker enables precision stratification and highlights an underserved, high-risk population requiring urgent development of alternative strategies like myeloid-targeted and interferon-directed therapies.

**Highlights:**

- Anti-RNP/Sm co-positivity defines a high-risk, flare-prone, myeloid-dominant SLE endotype
- RNP⁺Sm⁺ patients, especially Black patients, show profound refractoriness to first-line and B-cell depletion therapies
- Expanded intermediate monocytes with TLR4 upregulation and IFNA10-biased interferon signalling
- Hyperferritinaemia correlates with disease activity exclusively in RNP⁺Sm⁺ patients

## Introduction

Systemic lupus erythematosus (SLE) is a complex autoimmune disorder marked by adaptive and innate immune dysregulation. It disproportionately affects young women and ethnic minorities—especially Black and Asian populations—who experience higher rates of severe renal and cardiovascular complications, and for whom SLE is a leading cause of death during childbearing age (1–3). Despite focus on life-threatening organ involvement, outcomes have not improved: all-cause mortality remains nearly double, and deaths from renal, infectious, and cardiovascular causes are up to triple the general population’s rates (4, 5). Between 15-20% of patients still die prematurely, with minimal reduction in end-stage renal disease for the past 20 years (6)—underscoring a continuing unmet need.

Therapeutic progress in SLE has been limited. Dozens of randomised controlled trials (RCTs) have failed to meet primary endpoints, and—unlike rheumatoid arthritis with numerous approved biologics—approvals in SLE are limited to belimumab (anti-BAFF) and anifrolumab (type-I interferon receptor blockade), alongside off-label rituximab (anti-CD20). Recently, the FDA also approved obinutuzumab, another anti-CD20 monoclonal antibody, for the treatment of lupus nephritis. Response rates to these agents remain <50% and a modest ∼15-20% placebo-adjusted improvement; with newer anti-CD20 agents like obinutuzumab performing similarly (7–11). Attempts to combine B-cells therapies, such as sequential use of belimumab-rituximab to counter BAFF surges, have also shown limited benefit (12–14). Emerging and promising approaches such as B-cell-directed CAR-T offer “immune-reset” potential but face major barriers: high cost, complex delivery, profound immunosuppression, adverse effects, and lack of comparator data (15, 16).

It is plausible to suggest that the limited success of current therapies stems from the disease’s underlying immunologic heterogeneity (17–19). Multiple blood-based multi-omics studies integrating transcriptomics, proteomics and soluble mediators converge on at least two broad immune endotypes: a plasma cell/B-cell-skewed subtype and an innate subtype (18, 20–24). To facilitate the clinical translation of these endotypes using routinely collected serologic data, we adopted an unbiased, unsupervised analytical strategy integrating data from clinical trials (BEAT-Lupus, CALIBRATE, ACCESS) (12, 13, 25) and 2 observational cohorts. This approach revealed that patients with co-positivity for anti-RNP and anti-Sm antibodies (RNP⁺Sm⁺) represent a distinct subgroup. We therefore undertook a systematic characterisation of this endotype, combining clinical, immunophenotyping, cytokine profiling, Olink proteomics, HuProt autoantigen arrays, bulk RNA-seq, and pathway enrichment analyses. The following results describe the molecular, cellular, and clinical features of the RNP⁺Sm⁺ group, their associations with disease activity and organ involvement, and their relevance to clinical response across different therapeutic strategies.

## Results

The datasets used, and the rationale for their inclusion, are summarised in Supplementary Dataset Section 1 and Supplementary Table 1. Briefly, baseline data of BEAT-Lupus (active SLE refractory to first-line therapy) served as the primary discovery cohort because of its comprehensive and enriched clinical, laboratory and immunological profiling. Baseline values of CALIBRATE (refractory lupus nephritis) and ACCESS (lupus nephritis) were used as independent trial cohorts to assess treatment effects and validate key findings in renal-enriched populations. Two internal UCLH cohort provided external validation of incidence, clinical, laboratory and treatment-response associations in routine practice.

### RNP⁺Sm⁺: A Pro-inflammatory Endotype with Severe Disease

To identify autoantibody endotypes driving molecular heterogeneity, we mapped serological profiles using baseline samples from the BEAT-Lupus discovery cohort against integrated multi-omic dataset encompassing clinical, laboratory, cytokine, transcriptomic (including interferon/ISG signatures), and T- and B-cell immunophenotyping data. The serological panel comprised nine autoantibodies: conventional ENAs (Ro, La, RNP, Sm), IgG anti-cardiolipin, IgG anti-β2-glycoprotein I, IgG anti-dsDNA, and IgA1/IgA2 anti-dsDNA antibodies. IgA autoantibodies were included specifically based on our prior work demonstrating their relevance in SLE (26, 27). By integrating multi-omic data using MOFA+ (Multi-Omics Factor Analysis v2) (28), we defined the principal axes of biological heterogeneity in SLE. We then employed a distance-based BEST (BIO-ENV Stepwise Selection) algorithm (29) to determine which autoantibody combination most faithfully reproduced this molecular structure, thereby linking serologic classification to underlying biology.

The optimal subset emerged as co-positivity for anti-RNP and anti-Sm (RNP⁺Sm⁺). This signature showed a strong correspondence with the MOFA-derived molecular distance structure (Mantel r = 0.71; Spearman method; 9,999 permutations; nominal p = 0.0032; Figure 1A). This finding was robust to sensitivity analyses including only routinely measured lupus related autoantibodies (i.e. excluding IgA1/IgA2 anti-dsDNA antibodies) (Supplementary Figure 1A) and was validated across all three independent cohorts (CALIBRATE, ACCESS, UCLH) using the same MOFA–BEST framework, despite variable multi-omic depth (Supplementary Table 1 and Supplementary Figure 1B–D).

**Figure 1.**
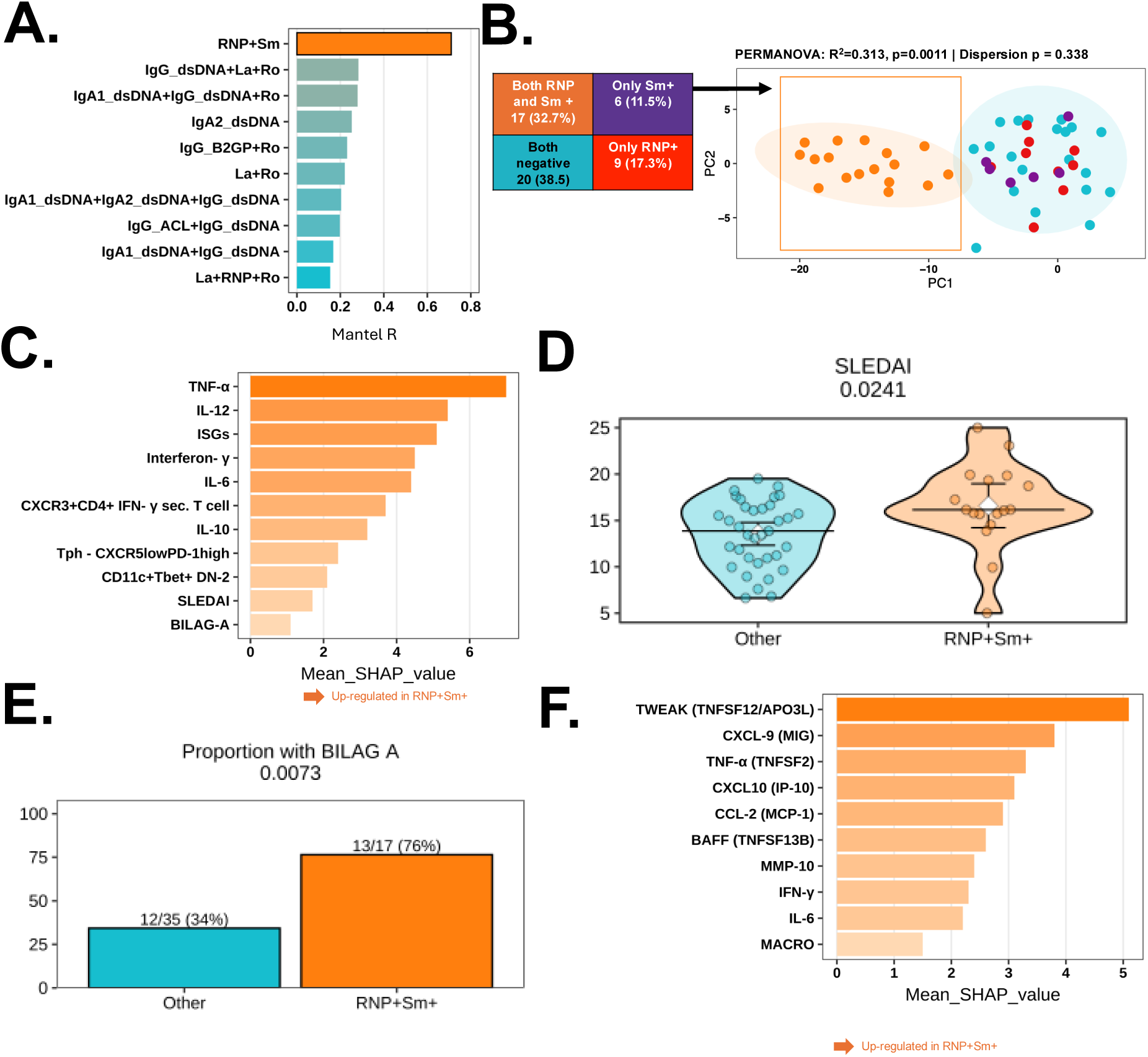
**A-F:** Immune clustering of lupus patients based on autoantibody profiles. BEAT-lupus trial baseline data (n=52): **A.** Multi-Omics Factor Analysis (MOFA) – Best Subset (BEST) serological subset analysis. Bars show the Mantel correlation (Spearman method) between serology-based dissimilarity (Jaccard distance on antibody positivity patterns for nine specificities: Sm, RNP, Ro, La, IgG anti-cardiolipin, IgG anti-β2-glycoprotein I, IgG anti-dsDNA, IgA1 anti-dsDNA, and IgA2 anti-dsDNA) and the global MOFA-derived dissimilarity (Euclidean distance across latent factors integrating clinical, cytokine, transcriptomic, interferon/ISG, and T- and B-cell immunophenotyping data). **B.** Ordination of the global biological distance matrix coloured by RNP⁺Sm⁺ status. Each point represents an individual patient; RNP⁺Sm⁺ patients are compared with those single-positive for either antibody or negative for both (“other” group). Group separation was assessed by permutational multivariate analysis of variance (PERMANOVA). **C.** Top 10 variables^$^ contributing to the separation in B, selected by a Random Forest (RF) classification algorithm and ranked by mean SHAP value, which quantifies each feature’s average impact on the model output. Pairwise comparisons of disease activity scores – **D.** SLEDAI-2004, and **E.** BILAG between RNP⁺Sm⁺ patients and the remaining patients (‘Other’). *p*-values for continuous variables were calculated using a linear regression model adjusted for age and ethnicity-Black vs. non-Black (unless otherwise indicated). Categorical variables were analysed using Fisher’s exact test. Black horizontal lines represent the median, white diamonds indicate the mean, and vertical lines denote 95% confidence intervals. **<u>Olink proteomics</u>** *<u>(Internal cohort – RNP⁺Sm⁺ n = 15, Other, n = 15):</u>* **D.** Bar plot showing the top differentially expressed plasma proteins (log₂ fold changes) in RNP⁺Sm⁺ versus others, as identified by RF and ranked by mean SHAP value. Fold changes are derived from normalised protein expression (NPX) values on a log₂ scale. *^$^ IgG_ACL = IgG anti-cardiolipin antibody, IgG_B2GP = IgG anti-beta2-glycoprotein, TNF-α = serum Tumour necrosis factor-α, IL-12 = serum Interleukin-12, IFN-γ = serum interferon gamma, CD11c⁺T-bet⁺ DN2 = CD11c⁺T-bet⁺ double negative (CD27^neg^IgD^neg^) memory B cells, IL-6 = serum Interleukin-6, Tph CXCR5^low^ PD-1^high^ - CXCR5^low^PD-1^high^CD3^+^CD4^+^ T-peripheral helper cells, IL-10 = serum Interleukin-10, SLEDAI = SLE Disease Activity Index 2000, CXCR3⁺ IFN-γ^+^ T cell = CXCR3⁺ IFN-γ-producing Th1 cells, BILAG = British Isles Lupus Assessment Group revised in 2004. TWEAK (TNF superfamily member 12; TNFSF12; also known as APO3 ligand), CXCL9 (C-X-C motif chemokine ligand 9; also known as monokine induced by gamma interferon [MIG]), TNF-α (Tumor necrosis factor alpha; TNF; also known as TNF superfamily member 2 [TNFSF2]), MACRO (Macrophage receptor MARCO; also known as SCARA2), M-CSF1 (Macrophage colony-stimulating factor 1; CSF1),CXCL10 (C-X-C motif chemokine ligand 10; also known as interferon gamma-induced protein 10 [IP-10]), CCL2 (C-C motif chemokine ligand 2; also known as monocyte chemoattractant protein-1 [MCP-1], MMP10 (Matrix metallopeptidase 10; also known as stromelysin-2), IFN-γ (Interferon gamma, BAFF (B-cell activating factor; also known as TNF superfamily member 13B [TNFSF13B])*

Permutational multivariate analysis of variance (PERMANOVA) confirmed that RNP⁺Sm⁺ status explained a substantial proportion of variance in the MOFA factor (R² = 0.31, P = 0.0011; 9,999 permutations; Figure 1B), with no evidence of unequal multivariate dispersion between groups (betadisper P = 0.34). This endotype accounted for 33% (17/52) of patients in the discovery cohort. Although all RNP⁺Sm⁺ patients were also IgG anti-dsDNA–positive (Supplementary Figure 2A), neither IgG anti-dsDNA titres nor conventional disease activity marker C3 levels differed between RNP⁺Sm⁺ and other patients across the four cohorts (Supplementary Figure 2B–E), indicating that other markers of disease activity are not significantly elevated in this endotype. Sensitivity analyses using PERMANOVA with predefined serological endotypes (IgG anti-dsDNA alone or combined with individual ENAs or antiphospholipid antibodies) showed these endotypes explained substantially less variance and formed less distinct clusters than RNP⁺Sm⁺ (Supplementary Table 2).

Random forest analysis identified key features differentiating this subgroup, dominated by inflammatory cytokines (Figure 1C). RNP⁺Sm⁺ patients had significantly higher disease activity, reflected by elevated SLEDAI scores (p=0.0241) and increased BILAG-2004 A manifestations (p=0.0073) (Figure 1D-E). To ensure phenotypic differences were not driven by overall disease activity score, all subsequent comparisons of continuous variables were adjusted for baseline SLEDAI-2K scores. Immunophenotyping revealed enriched pro-inflammatory cell subsets: peripheral T helper (CXCR5low PD-1high) cells (p=0.0167), CD11c⁺T-bet⁺ DN2 memory B-cells (p=0.0246), and CXCR3⁺ IFN-γ-producing Th1 cells (p=0.0347) (Supplementary Figure 1A–C). In addition, several serum cytokines including TNF-α (p=0.0062), IL-12 (p=0.0151), IL-6 (p=0.0224), IFN-γ (p=0.0177), and IL-10 (p=0.0077) were significantly elevated (Supplementary Figure 1D–H). RNP⁺Sm⁺ patients also revealed greater interferon-stimulated gene (ISGs) expression (p=0.0244) (Supplementary Figure 3I).

To isolate the synergistic impact of co-positivity, we performed three-way comparisons (RNP⁺Sm⁺ vs. single-positive vs. double-negative). These demonstrated that single-positive patients displayed phenotypes indistinguishable from double-negative controls (Supplementary Figure 4). Plasma inflammatory proteomics further corroborated this heightened inflammatory state, showing elevated TWEAK, CXCL-9, TNF-α, and CXCL10 in this endotype (Figure 1F).

Finally, we validated these findings using data from the CALIBRATE trial. Although CALIBRATE predominantly enrolled patients with lupus nephritis—representing a clinically distinct population from BEAT-Lupus—the results were consistent. At baseline RNP⁺Sm⁺ patients in CALIBRATE similarly exhibited higher SLEDAI scores (p = 0.0008), increased DN2 B-cell frequencies (p = 0.0049), and elevated serum IL-6 concentrations (p = 0.0423) compared to the rest of the cohort (Supplementary Figures 5A-D).

### RNP⁺Sm⁺: Predominant in Black patients with nephritis/vasculitis/enteritis and high flare burden with raised Ferritin

RNP⁺Sm⁺ positivity was significantly enriched among treatment-refractory patients requiring biologic therapy or cyclophosphamide compared to non-refractory patients (p=0.0002; Figure 2A). The RNP⁺Sm⁺ endotype was also more frequent in Black patients (p = 0.0002; Figure 2B), while gender distribution remained comparable across groups.

**Figure 2.**
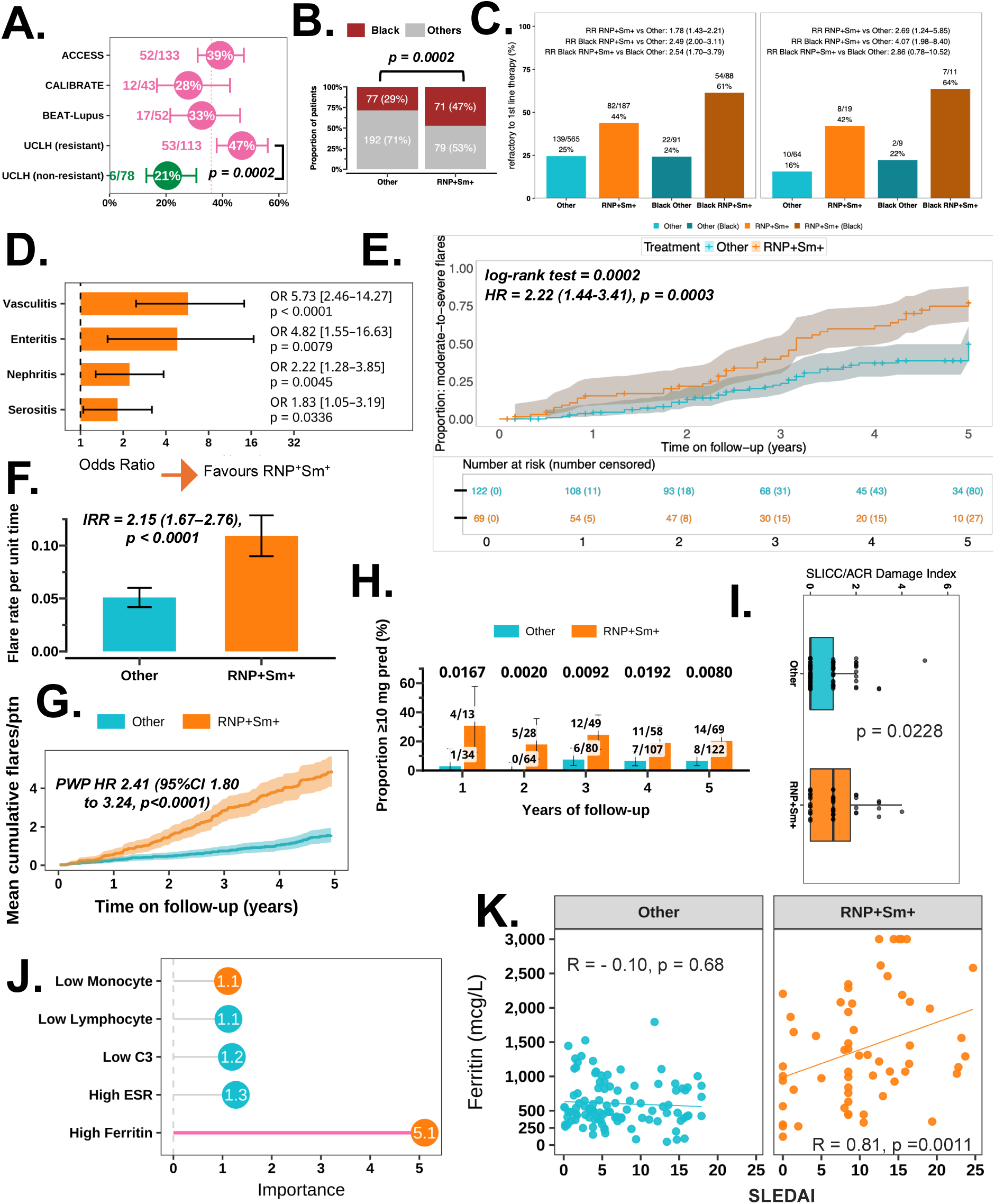
**A-K:** Phenotypic characterisation of the RNP⁺Sm⁺ endotype (positive for both anti-RNP and anti-Sm antibodies). **A.** Proportion of patients with RNP⁺Sm⁺ positivity across three randomised controlled trials (BEAT-Lupus, CALIBRATE, ACCESS) and one internal observational cohort (UCLH) *^$^*. Patients in the RCTs represent treatment-refractory disease (requiring biologic therapy or cyclophosphamide). The UCLH cohort comprised both treatment-refractory and non-refractory patients. p-values comparing refractory versus non-refractory groups were calculated using Fisher’s exact test. The vertical dashed line indicates the pooled mean proportion of RNP⁺Sm⁺ patients. **B.** Proportion of Black patients (self-identified or reported as Black ancestry) versus other ethnic backgrounds among RNP⁺Sm⁺ cases, based on pooled data from all four cohorts (comparison by Fisher’s exact test). **C.** Proportion of refractory patients requiring escalation to biologics or cyclophosphamide. Two independent cohorts (UCLH cohort-left and CUH cohort-right) were analysed and stratified by co-positivity for anti-RNP and anti-Sm antibodies (RNP⁺Sm⁺) versus any other autoantibody combination (“Other”), and by ethnicity (Black vs non-Black). Risk ratios (RR) with 95% confidence intervals compare refractory risk among the groups. **D.** Organ involvement in RNP⁺Sm⁺ patients analysed using penalised multivariate logistic regression adjusted for age, and ethnicity (Black vs. Non-Black). Organ variables were selected via a random forest feature selection algorithm. This analysis combined data from BEAT-Lupus, CALIBRATE, and the observational cohort (excluding ACCESS due to unavailable detailed BILAG-2004 scores). Bar charts display adjusted odds ratios (OR) and 95% confidence intervals. Vasculitis: RNP⁺Sm⁺ *n*=19/98, Other *n*=9/188; Enteritis: RNP⁺Sm⁺ *n*=9/98, Other *n*=5/188; Nephritis: RNP⁺Sm⁺ *n*=41/98, Other *n*=48/188; Serositis: RNP⁺Sm⁺ *n*=38/98, Other *n*=49/188. **E.** Kaplan–Meier analysis of time to first moderate-to-severe flare (BILAG-A or 2B) in the observational cohort. Hazard ratios were estimated using Cox proportional hazards regression adjusted for age, ethnicity, baseline SLEDAI, and medication dose. **F.** Observed rates of moderate-to-severe flare during follow-up. Incidence rate ratios (IRR) and 95% confidence intervals were calculated using Poisson regression adjusted for age, ethnicity, baseline SLEDAI, and medication dose (observational cohort). **G.** Recurrent moderate-to-severe flares beyond the first event (observational cohort). Mean cumulative function curves show the average number of flares per patient over follow-up (shaded areas indicate 95% CIs). The hazard ratio (HR) for recurrent flares was estimated using a Prentice–Williams–Peterson (PWP) total-time Cox model, stratified by event order with robust standard errors, and adjusted for age, ethnicity, baseline SLEDAI, and medication dose. **H.** Proportion of patients receiving prednisolone ≥10 mg/day during observation. p-values at each time point were calculated using Fisher’s exact test. Patient numbers are indicated above each bar. **I.** Comparison of SLICC/ACR Damage Index scores at 5 years, p-value adjusted for age, ethnicity, baseline SLEDAI, and medication dose. **J.** Conventional laboratory markers at the time of flare in RNP⁺Sm⁺ (*n*=42) versus other patients (*n*=42), analysed using a random forest model. The top five variables ranked by importance score (Mean Decrease Gini) are displayed. Orange bars represent variables associated with flares in RNP⁺Sm⁺ patients; sky blue bars represent variables associated with other patients. The pink line indicates the sole variable (Ferritin) retaining significance following feature selection by the Boruta algorithm. **K.** Scatter plots showing the relationship between serum ferritin and disease activity (SLEDAI score) in patients from the combined UCLH and CUH cohorts. Analyses are stratified by autoantibody status: (Left) Other SLE endotypes (*n* = 102) and (Right) RNP⁺Sm⁺ endotype (positive for both anti-RNP and anti-Sm antibodies; *n* = 59). Spearman’s rank correlation coefficients (rho) and p-values are displayed for each group. *^$^ Clinical Trials - BEAT-Lupus trial (RNP⁺Sm⁺, n = 17; Other, n = 35), CALIBRATE trial (RNP⁺Sm⁺, n = 12; Other, n = 31), and ACCESS trial (RNP⁺Sm⁺, n = 52; Other, n = 85); Observational Internal cohort comprised new patients seen at the lupus clinic between June 2019 and June 2024 (RNP⁺Sm⁺, n = 69; Other, n = 122)*.

In our cohort, 44% of patients with RNP⁺Sm⁺ (82/187) failed to achieve an adequate response to first-line conventional immunosuppressive therapy, necessitating escalation to biologics or cyclophosphamide, compared to 25% of those who were not co-positive for anti-RNP and anti-Sm antibodies (139/565; risk ratio/RR 1.78, 95%CI 1.43–2.21) (Figure 2C). Non-response rates rose strikingly to 61% (54/88) among Black RNP⁺Sm⁺ patients (RR 2.49 vs. the other patients, 95%CI 2.00–3.11). Crucially, another independent cohort validated this refractory phenotype in mSLE.

To identify associated clinical manifestations, random forest analysis was performed across three independent datasets: BEAT-Lupus, lupus nephritis RCTs-CALIBRATE, and an observational cohort, which revealed systemic vasculitis, lupus nephritis, enteritis, and serositis as positively associated with RNP⁺Sm⁺. Multivariate penalised logistic regression confirmed these associations, with systemic vasculitis demonstrating the strongest relationship (penalised OR/pOR: 5.73, 95%CI: 2.46–14.27; p<0.0001), followed by enteritis (pOR: 4.82, 95%CI: 1.55–16.63; p=0.0079) and nephritis (pOR: 2.22, 95%CI: 1.28–3.85; p=0.0045) (Figure 2D; Supplementary Figure 6).

Time-to-event analysis of longitudinal observational data revealed that RNP⁺Sm⁺ patients experienced significantly shorter time to first moderate-to-severe flare, defined as BILAG-A or BILAG-2B events (log-rank p=0.0002; Figure 2E). After adjusting for age, hydroxychloroquine, immunosuppressants, and prednisolone dose, the hazard ratio for severe flare in RNP⁺Sm⁺ patients was 2.22 (95%CI: 1.44–3.41; p=0.0003). The incidence rate ratio for moderate-to-severe flares was 2.15 (95%CI: 1.67–2.76; p<0.0001; Figure 2F). Recurrent-event analysis using a Prentice-Williams-Peterson total-time Cox model, stratified by event order with identical covariate adjustments, confirmed persistently elevated flare risk over five years (HR: 2.41, 95%CI: 1.80–3.24; p<0.0001; Figure 2G). Consistent with this moderate-to-severe disease activity, RNP⁺Sm⁺ patients required prednisolone ≥10 mg/day more frequently during follow-up (Figure 2H); thereby showed increased accrual of ACR-EULAR SLICC defined damage (p=0.0228, Figure 2I).

### Hyperferritinaemia as a Signature of the RNP⁺Sm⁺ Endotype

At the onset of disease flare, markedly elevated serum ferritin was the most distinct feature of this endotype (p < 0.0001), reflecting underlying macrophage activation (Figure 2J, Supplementary Figure 7A–B). Ferritin levels correlated strongly with key pro-inflammatory cytokines, including TNF-α (r = 0.81, p = 0.0003), IL-6 (r = 0.78, p = 0.0015), and IFN-γ (r = 0.71, p = 0.0034), defining a cytokine–ferritin network characteristic of myeloid and type I/II interferon-driven inflammation (Supplementary Figure 7C). In contrast, ferritin showed no significant correlation with CRP (r = 0.15, p = 0.32).

The association between hyperferritinaemia and active disease was subsequently validated in the independent CUH cohort (Supplementary Figure 7D). Notably, in a combined analysis of both cohorts, serum ferritin levels exhibited a strong correlation with SLEDAI scores exclusively within the RNP⁺Sm⁺ group (r=0.81, p=0.0011; Figure 2K).

### Poor Response to B Cell Depletion Therapy

The RNP⁺Sm⁺ endotype demonstrated poor clinical response to rituximab across three clinically distinct cohorts. Meta-analysis revealed a 57% relative reduction in response (RR=0.43; 95%CI 0.29–0.58; p=0.0112; Figure 3A), with a number needed to harm of 2.33 (95%CI 1.72–3.45). Pooling the three cohorts, a Bayesian hierarchical model estimated that RNP+Sm+ patients had a posterior mean absolute rituximab response rate 43.5 percentage points lower than other patients (95% credible interval 14.4 to 72.8 percentage points lower; Figure 3B). The posterior probability that this endotype truly has a poorer response was 99.2% (Pr = 0.992).

**Figure 3.**
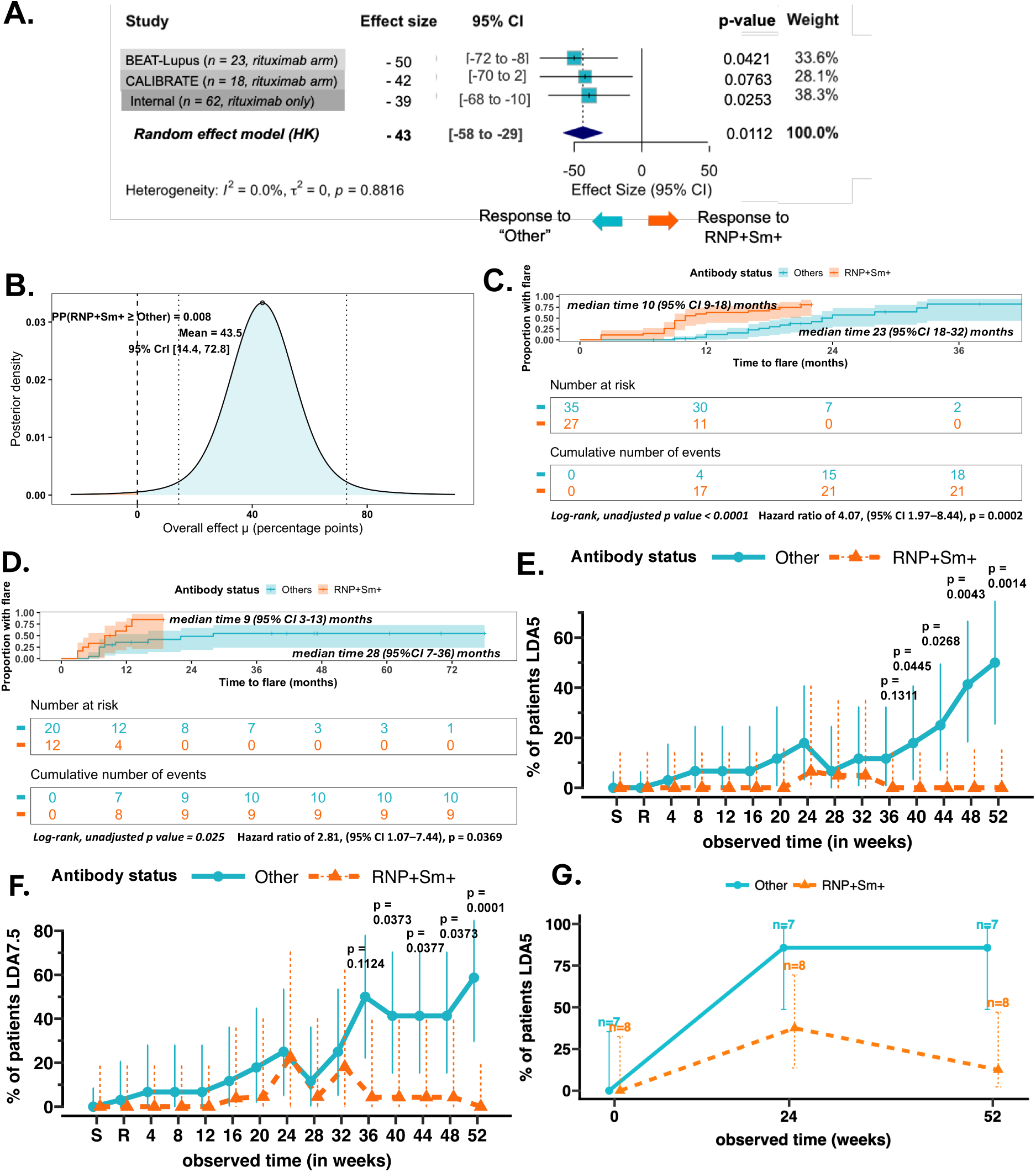
**A-G:** RNP⁺Sm⁺ (positive for both anti-RNP and anti-Sm antibodies) endotype is associated with poor response and higher flare rates following rituximab therapy. A. Clinical Response to Rituximab: The forest plot depicts response rates to rituximab in the BEAT-Lupus and CALIBRATE trials, and an internal observational cohort, evaluated at 52 weeks, 48 weeks, and 24 weeks, respectively. Patients in the observational cohort were treated with rituximab as part of standard NHS England care alongside other therapies. Effect sizes and 95% confidence intervals are shown for each cohort. The pooled effect size, shown as a diamond, was estimated using a random-effects meta-analysis model (Hartung-Knapp adjustment). Heterogeneity was assessed using *I^2^*. p-values shown here calculated by logistic regression and adjusted for age, ethnicity (Black vs Non-Black), and lupus-nephritis. **B.** Bayesian posterior for the pooled absolute difference in response. Curve shows the posterior density of the overall mean effect (μ) from a Normal–Normal random-effects meta-analysis (flat prior on μ; half-Cauchy(0, 0.5) prior on τ) across the 3 cohorts. Shaded curves are posteriors from a beta–binomial model with Jeffreys prior Beta(0.5,0.5) fitted to pooled responder/non-responder counts from the three studies. **C-D.** <u>Time to flare and flare rates in</u> <u>patients treated with rituximab</u>: **C.** Kaplan–Meier estimate of time to flare (moderate-to-severe: BILAG-2004 A or 2B) in the internal observational cohort. Adjusted flare rates were calculated using Cox proportional hazards regression, adjusting for age, ethnicity, medication dose and lupus-nephritis at baseline in RNP⁺Sm⁺ patients over the observation period. Log-rank p values, adjusted hazard ratios, and median times to event are shown. **D.** Re-analysis of previously published data, showing Kaplan–Meier curves for time to flare. Between-group differences were assessed using the log-rank test, with Cox proportional hazards regression used to calculate hazard ratios and p-values. **E-F.** <u>Proportion of patients achieving persistent</u> <u>low disease activity (LDA)</u>*^$^*on **E.** ≤5 mg/day prednisolone (LDA5) and **F.** ≤7.5 mg/day prednisolone (LDA7.5), stratified by RNP⁺Sm⁺ endotype versus all other patients in the rituximab arm (total n=26; RNP⁺Sm⁺ n=8; Other n=18) of the BEAT-Lupus trial. S = screening (6–8 weeks prior to randomisation); R = randomisation. **G.** Proportion of patient achieving LDA5 stratified by antibody status following obinutuzumab. Lines show proportions (95% CIs) of patients in LDA at each visit by group, estimated from a linear mixed-effects model of the arcsine–square-root–transformed proportion (random intercept for participant; fixed effects for group, time [visit as factor], and their interaction). Estimated marginal means were back-transformed to the proportion scale and plotted as percentages. p-values (where shown) were calculated using Tukey-adjusted pairwise contrasts between groups at each visit, with adjustment for age, ethnicity, baseline medication dose, and lupus nephritis status. *^$^Persistent LDA is defined as no BILAG-2004 Grade A or B scores in any organ system at 2 consecutive visits 4 weeks apart (all organ systems scored as C, D, or E), with prednisone dose ≤5 mg/day (LDA5) or ≤7.5 mg/day (LDA7.5) at both visits, and stable immunosuppressant and hydroxychloroquine dose for the preceding 12 weeks*.

Time-to-event analyses in both the observational cohort and re-analysed previously published datasets (30) consistently demonstrated earlier moderate-to-severe disease flares following rituximab treatment among RNP⁺Sm⁺ patients (log-rank p<0.0001 and p=0.025, respectively; Figure 3C–D). Critically, no RNP⁺Sm⁺ patients treated with rituximab achieved BILAG persistent low disease activity (LDA) at 52 weeks, compared to half the non-RNP⁺Sm⁺ patients achieving BILAG persistent LDA with prednisolone ≤5 mg/day (50%, 95%CI: 24–76%; p=0.0014) or ≤7.5 mg/day (58%, 95%CI: 29–88%; p=0.0001) (Figure 3E-F). Similarly, the newer B-cell-depleting agent obinutuzumab demonstrated reduced efficacy, with lower LDA at 52 weeks (p=0.0101, Figure 3G).

Although adding belimumab after rituximab in BEAT-Lupus and CALIBRATE did not result in inferior response rates among RNP⁺Sm⁺ patients compared with other serological groups, neither belimumab (sequential or monotherapy) nor cyclophosphamide-based regimens yielded better outcomes in this endotype, despite its myeloid-dominant and BAFF-enriched immunological profile (Supplementary Figures 8A–B and Supplementary Figures 9A–C).

To investigate mechanisms underlying poor response to rituximab, we further analysed longitudinal CD19⁺ B-cell counts in the BEAT-Lupus trial. RNP⁺Sm⁺ patients treated with rituximab demonstrated a similar degree of B cell depletion but significantly earlier B-cell repopulation at 24 weeks (p<0.0001), which persisted through 52 weeks (p<0.0001) compared to other patients (Figure 4A). While no differences emerged in the belimumab arm (Figure 4B), RNP⁺Sm⁺ patients demonstrated consistently shorter time to CD19⁺ repopulation across two independent rituximab-treated cohorts: BEAT-Lupus (24 versus 52 weeks; log-rank p = 0.0065; Figure 4C) and an observational cohort (28 versus 52 weeks; log-rank p = 0.0135; Supplementary Figure 10A). Similar to rituximab, RNP⁺Sm⁺ patients also showed earlier B-cell repopulation after the newer-generation B-cell-depleting agent, obinutuzumab (p=0.0199), from our internal observational cohort (Figure 4D). In the CALIBRATE trial where both arms received cyclophosphamide with rituximab, CD19⁺ B-cell repopulation rates were slower in the belimumab arm compared to placebo, but did not differ between RNP⁺Sm⁺ and other serological subgroups (Supplementary Figure 10B-C).

**Figure 4.**
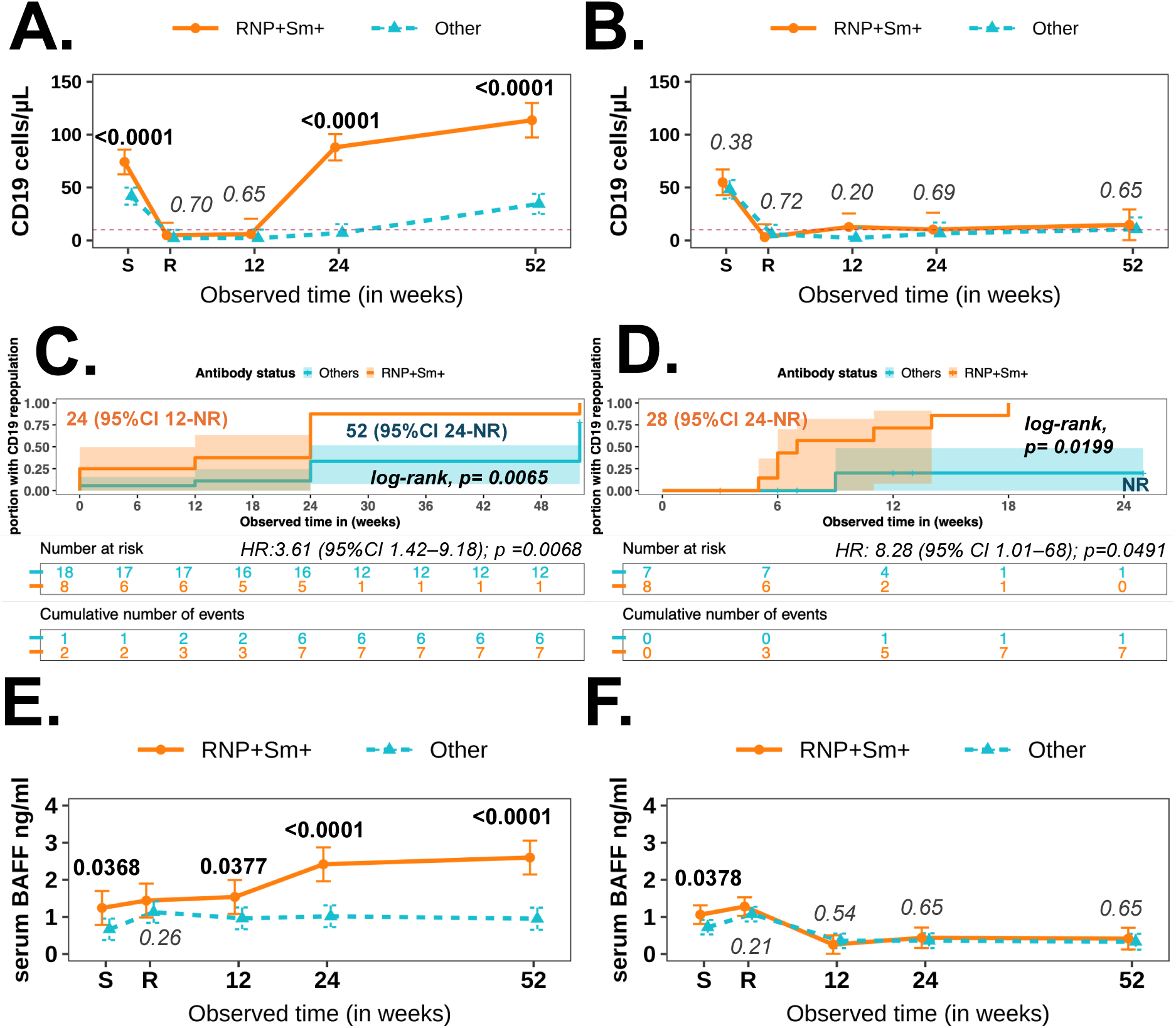
**A–F:** Repopulation of CD19⁺ B-cell counts in the RNP⁺Sm⁺ endotype (positive for both anti-RNP and anti-Sm) treated with rituximab-only in the BEAT-Lupus trial correlated with temporal rise in serum BAFF (B-cell activating factor). Changes in CD19⁺ B cells in **A.** placebo–rituximab arm and **B.** belimumab–rituximab arm, from screening (S), randomisation (R) to week 52. Horizontal dashed line marks 10 cells/µL (B-cell depletion threshold). **C.** Kaplan–Meier curves for time to CD19⁺ B-cell repopulation; Cox regression adjusted for age, ethnicity, nephritis, and medication dose. Log-rank p values, adjusted hazard ratios, and median times to event are shown. NR = median value not reached. **D.** Internal observational cohort-patients treated with obinutuzumab: Kaplan–Meier curves for time to CD19⁺ B-cell repopulation; Cox regression adjusted as above. Serum BAFF changes in **E.** placebo–rituximab arm and **F.** belimumab–rituximab arm, from screening (S) to week 52 in BEAT-Lupus. Lines show geometric means (95% CIs) by group over time, estimated from a mixed-effects model with a log link (random intercept for participant; fixed effects for group, time, and their interaction). Estimated marginal means were back-transformed to the original scale. p-values were calculated using Tukey-adjusted pairwise contrasts between groups at each visit, with adjustment for age, ethnicity, baseline medication dose, and lupus nephritis status.

Given BAFF’s role in B-cell reconstitution following rituximab, we examined longitudinal serum BAFF levels. In BEAT-Lupus baseline serum BAFF was elevated in RNP⁺Sm⁺ patients and rose more rapidly from 12 weeks onwards in the rituximab only group (p=0.0377 at 12 weeks; p<0.0001 at 24 and 52 weeks), temporally correlating with accelerated CD-19 repopulation (Figure 4E). This accelerated increase in serum BAFF did not occur in the belimumab arm, which neutralises soluble BAFF (Figure 4F). Similarly, in CALIBRATE, where cyclophosphamide was combined with rituximab, BAFF levels did not differ between the RNP^+^Sm^+^ endotype and the other subgroup (Supplementary Figure 10D-E), paralleling the absence of differential B cell repopulation.

To further investigate temporal changes in ISGs and cytokines that were elevated at baseline in the BEAT-Lupus trial, we performed longitudinal analyses which showed that RNP⁺Sm⁺ patients maintained a persistently heightened ISG and cytokine profile in the rituximab arm, whereas belimumab attenuated several of these mediators, including IL-6 (p = 0.0129) and IL-12 (p = 0.0085) (Supplementary Figure 11). This sustained interferon–myeloid cytokine milieu in the rituximab arm provides further mechanistic support for the lack of clinical response to B-cell depletion in this endotype.

### RNP⁺Sm⁺ endotype as a proinflammatory, myeloid-skewed phenotype

M-scores for immune-related gene modules, computed from bulk RNA-seq data (BEAT-Lupus, ACCESS, and published datasets) and mapped to 606 standardised modules, revealed distinct pathway signatures with significant upregulation of interferon-related, monocyte/inflammation-related, and neutrophil-related pathways, whereas B-cell signatures were preferentially elevated in non-RNP⁺Sm⁺ patients (Figure 5A). CIBERSORTx deconvolution of whole-blood transcriptomes, using reference immune-cell expression signatures, confirmed these findings by revealing significantly higher inferred cell-type fractions of monocytes (p=0.0009), proinflammatory M1 macrophages (p<0.0001), and neutrophils (p=0.0048) in RNP⁺Sm⁺ patients (Figure 5B), whereas plasma cells (p=0.0018) and memory B cells (p=0.0055) were relatively enriched in non–RNP⁺Sm⁺ patients.

**Figure 5.**
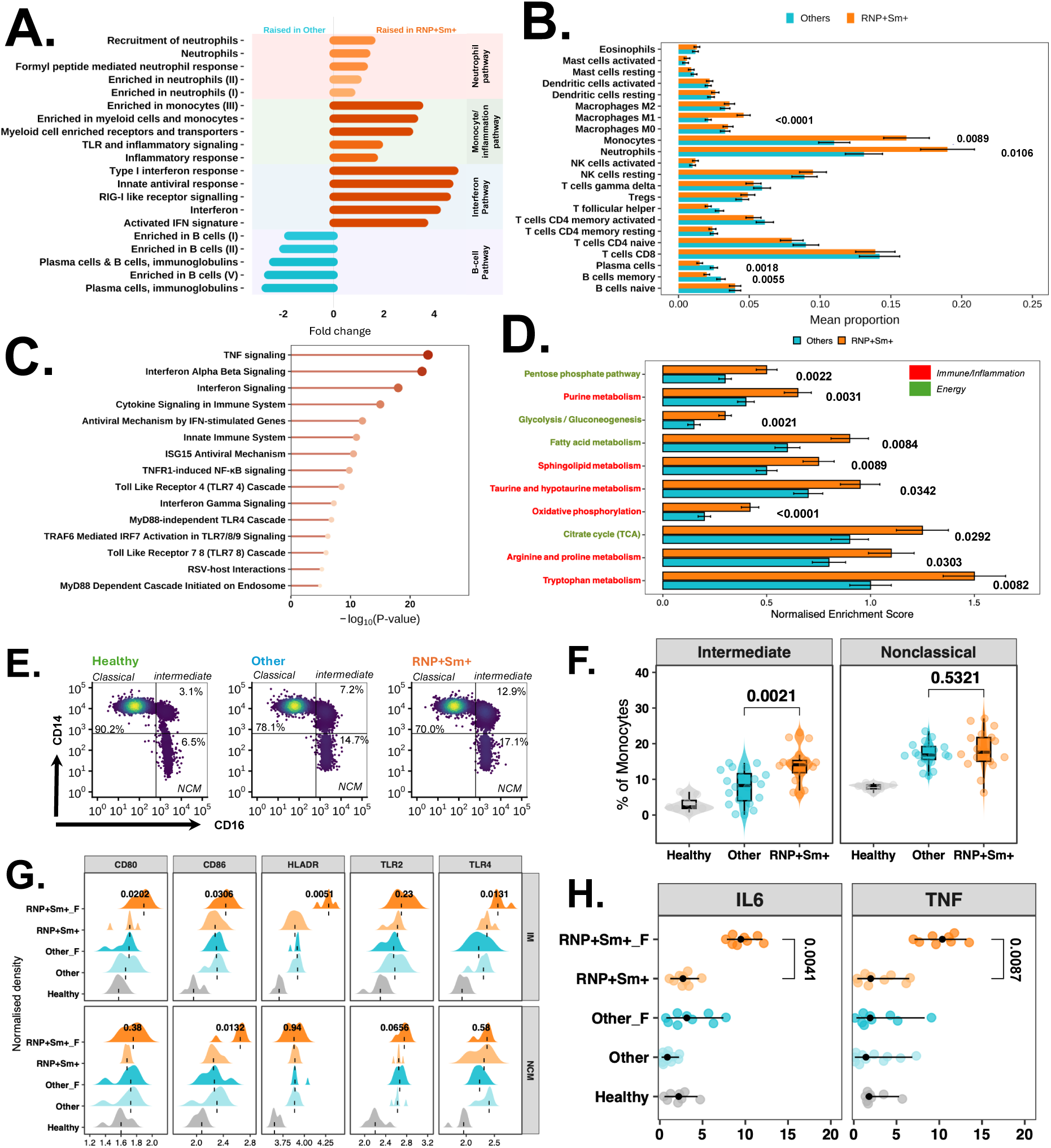
**A-G:** Transcriptomic and metabolic signatures along with expansion and activation of intermediate monocytes (IMs) in lupus patients with anti-RNP and anti-Sm antibodies (RNP⁺Sm⁺) compared to other lupus patients*^$^*. A. Bar charts showing module-level expression (M-scores) for immune-related blood transcriptional modules comparing RNP⁺Sm⁺ patients with other lupus patients. For each individual, gene expression was standardised to z-scores relative to healthy controls, and module M-scores were calculated as the mean z-score of all genes within each module. Displayed modules are grouped into four functional categories: B cell–related modules, interferon-related modules, monocyte/inflammation-related modules, and neutrophil-related modules. **B.** Immune cell composition estimated from bulk RNA-seq data using CIBERSORTx with the LM22 signature matrix, representing 22 immune cell subsets. Bars show mean proportions with error bars representing standard deviation. p-values were calculated using Mann–Whitney U test and corrected for multiple testing using the Benjamini-Hochberg method. **C.** Reactome pathway enrichment analysis of upregulated differentially expressed genes (DEGs) in the RNP⁺Sm⁺ endotype, based on bulk RNA-seq data. DEGs were defined using an FDR < 0.05 and |log₂FC| > 0.5 when comparing RNP⁺Sm⁺ patients with other lupus patients. **D.** KEGG pathway enrichment for metabolism-associated DEGs: Bars represent normalised enrichment scores (NES) for RNP⁺Sm⁺ versus other patients, with pathway names color-coded by functional category. **E.** Representative flow cytometry plots showing monocyte subset gating based on CD14 and CD16 expression in healthy controls, RNP⁺Sm⁺, and other SLE patients. Monocyte subsets are defined as classical (CD14⁺⁺CD16⁻), intermediate (CD14⁺⁺CD16⁺), and non-classical/NCM (CD14⁺CD16⁺⁺). **F.** Frequencies of CD16⁺ monocyte subsets (intermediate and non-classical) among total peripheral blood monocytes. Violin plots show individual data points with overlaid medians and interquartile ranges. P values are derived from linear regression models of logit-transformed subset frequencies, including group, age, and ethnicity as covariates, with Benjamini–Hochberg correction for multiple testing. Group sizes: RNP⁺Sm⁺, n = 17; Other SLE, n = 20; Healthy, n = 8. **G.** Expression of activation and innate immune-sensing markers on intermediate (IM, upper panels) and non-classical (NCM, lower panels) monocytes. Density ridge plots display marker expression distributions across groups: Healthy controls, Other SLE (non-RNP⁺Sm⁺ patients in remission, n = 11), Other_F (non-RNP⁺Sm⁺ patients with flare, n = 9), RNP⁺Sm⁺ (RNP⁺Sm⁺ patients in remission, n = 9), and RNP⁺Sm⁺_F (RNP⁺Sm⁺ patients with flare, n = 8). X-axes show fluorescence intensity on a log₁₀ scale. Ridge heights represent normalised kernel density estimates; black dashed vertical lines indicate medians. Where reported, P values are obtained from linear regression models of log-transformed fluorescence intensity, adjusted for age and ethnicity, with Benjamini–Hochberg correction for multiple testing. **H.** Intracellular cytokine staining of IM. Black dots indicate medians and lines the interquartile ranges. P values are from linear regression models of log-transformed cytokine signal comparing RNP⁺Sm⁺ and RNP⁺Sm⁺_F, adjusted for age and ethnicity, with Benjamini–Hochberg correction for multiple testing. *^$^ Participants were drawn from three cohorts: (i) the BEAT-Lupus clinical trial (RNP⁺Sm⁺, n = 11; Other, n = 22); (ii) the ACCESS trial (RNP⁺Sm⁺, n = 52; Other, n = 85), and (iii) publicly available data (GSE72326) (RNP⁺Sm⁺, n = 30; Other, n = 127)*.

Reactome pathway enrichment analysis of upregulated differentially expressed genes (DEGs; FDR<0.05, log₂FC>0.5) revealed significant enrichment of TNF and IFN signalling, TNF receptor-induced NF-κB activation, MyD88-dependent signalling, and Toll-like receptor cascades (Figure 5C). KEGG pathway analysis demonstrated broad metabolic reprogramming in RNP⁺Sm⁺ patients, with upregulation of amino acid catabolism (tryptophan, arginine/proline, and taurine/hypotaurine), lipid metabolism (fatty acid and sphingolipid biosynthesis), central carbon metabolism (glycolysis, pentose phosphate pathway, TCA cycle, and oxidative phosphorylation), and nucleotide synthesis (purine metabolism) compared with other patients (Figure 5D).

To validate the monocyte phenotype characterising this endotype, flow cytometric analysis of stored PBMCs from a subset of lupus patients demonstrated significant expansion of intermediate monocytes (CD14⁺⁺CD16⁺; IM) in RNP⁺Sm⁺ patients (p=0.0021; Figure 5E–F, Supplementary Figure 12). While antigen-presenting molecules (CD80, CD86, and HLA-DR) were elevated in both IM and non-classical monocytes (CD14⁺CD16⁺⁺; NCM) across all patients irrespective of activity status, the upregulation was only significant in IM from active/flared (BILAG-A or B) RNP⁺Sm⁺ patients, specifically for HLA-DR (p=0.0051) and CD80 (p=0.0202) (Figure 5G). Furthermore, flare-associated IM from RNP⁺Sm⁺ patients demonstrated enhanced inflammatory capacity, with increased TLR4 surface expression (p=0.0131) (Figure 5G) and higher frequencies of IL-6-(p=0.0041) and TNF-α-producing (p=0.0087) IM (Figure 5H).

### IgG autoreactivity predominated against spliceosomal and RNA-splicing–related proteins in the RNP⁺Sm⁺ endotype

Using HuProt proteome microarrays, we profiled both IgG and IgA autoantibody reactivity from BEAT-Lupus samples stratified according to anti-RNP anti-Sm co-positivity, and applied pre-ranked gene set enrichment analysis with MSigDB annotations (GO:CC, GO:BP, Reactome). IgG-associated autoreactivity was strongly enriched for spliceosome and RNA-processing components, consistent with the canonical co-targets of anti-RNP and anti-Sm autoantibodies (Figure 6A, Supplementary Figure 13A). By contrast, IgA responses did not show significant enrichment in these top spliceosome modules (Figure 6B, Supplementary Figure 13B). Normalised enrichment scores (RNP⁺Sm⁺ versus Other) confirmed that spliceosomal signatures were largely restricted to IgG (Supplementary Figure 13C).

**Figure 6.**
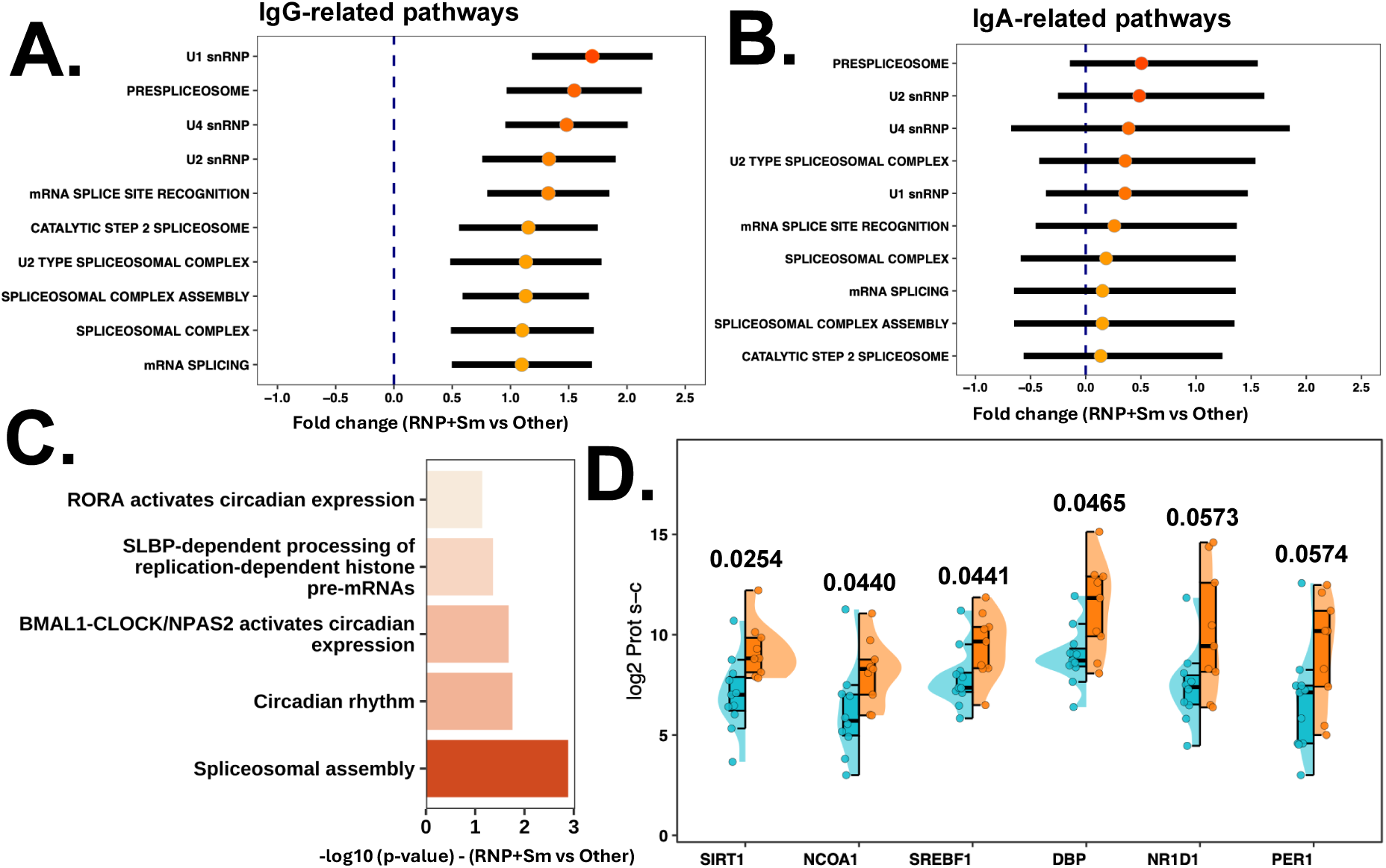
**A–D:** IgG-associated autoreactivity is enriched for spliceosome and circadian pathway–targeted modules in RNP⁺Sm⁺ patients (BEAT-Lupus cohort). Bar plots display **A.** IgG and **B.** IgA-associated autoreactivity scores for nine spliceosome-related modules, among the fifteen most significantly enriched functional sets identified by pre-ranked gene set enrichment (cameraPR) analysis using MSigDB annotations (GO:CC, GO:BP, Reactome). Bars show the mean difference in module scores between RNP⁺Sm⁺ (n=9) and other patients (n=11) (Δ = RNP⁺Sm⁺ – Other), with 95% confidence intervals, adjusted using corresponding healthy controls from each batch. **C.** Top Reactome functional pathways enriched among differentially targeted IgG autoantigens, based on a false discovery rate (FDR, Benjamini–Hochberg) < 0.05 and a log₂ fold change (log₂FC) > 0.5; bar height reflects statistical significance (–log₁₀[p-value]), with p-values indicated above each bar. **D.** Mean log₂ protein microarray signal intensities for IgG autoantibodies directed against top six core circadian rhythm regulators, comparing RNP⁺Sm⁺ double-positive patients to other lupus patients. Bars represent group means with 95% confidence intervals; adjusted (age and ethnicity) p-values from Wilcoxon rank-sum tests are shown above each comparison to indicate statistical significance.

Beyond spliceosome-associated pathways, enrichment analysis of differentially targeted IgG autoantigens between RNP⁺Sm⁺ and other patients revealed significant over-representation of circadian rhythm-related pathways, including circadian clock regulation, BMAL1–CLOCK/NPAS2-mediated transcription, and RORA-driven circadian expression (Figure 6C). This enrichment reflected autoreactivity against classical circadian regulators (NR1D1/REV-ERBα, PER1, DBP) alongside metabolic and epigenetic effectors (SIRT1, NCOA1, SREBF1) (Figure 6D). To identify genes with the greatest differential targeting, a composite ranking approach was applied, integrating effect size (Δ log₂ signal between RNP⁺Sm⁺ and non-RNP⁺Sm⁺ patients) and false discovery rate-adjusted Wilcoxon p-values. SIRT1 ranked highest (p=0.0254), followed by NCOA1 (p=0.0440) and SREBF1 (p=0.0441), indicating prominent IgG-mediated autoreactivity against circadian-metabolic regulators in RNP⁺Sm⁺ SLE.

To determine whether autoantibody reactivity exhibited a serological gradient, Jonckheere–Terpstra trend analyses were performed across RNP⁺Sm⁺ double-positive (Both⁺), single-positive (Either⁺), and seronegative (Neither) groups using spliceosome module scores. Most spliceosome modules demonstrated significant positive trends (Δ > 1, adjusted p<0.05; Supplementary Figure 14A), reflecting graded increases in autoreactivity with highest levels in RNP⁺Sm⁺ patients. In contrast, IgA-associated spliceosome modules showed overall lower module scores and no significant trends (Supplementary Figure 14B), confirming preferential IgG-mediated autoreactivity in this endotype. Paralleling the autoreactivity gradient, cytokine measurements showed ordered increases in IFN-α, IFN-γ, BAFF, IL-6, and TNF-α from Neither to Either⁺ to Both⁺ groups (Supplementary Figure 15A–F).

### IFNA10-Driven Interferon Signatures in RNP⁺Sm⁺ Lupus

Among RNP⁺Sm⁺ patients with available proteome microarray data, type I ISG module scores were significantly elevated (p=0.0093; Figure 7A), and were best predicted by IFNA10 (interferon-α10) mRNA expression (Figure 7B). Although total serum IFN-α concentrations were similar between groups (p=0.29; Figure 7C), IFNA10 mRNA was selectively increased in RNP⁺Sm⁺ individuals (p=0.0004; Figure 7D).

**Figure 7.**
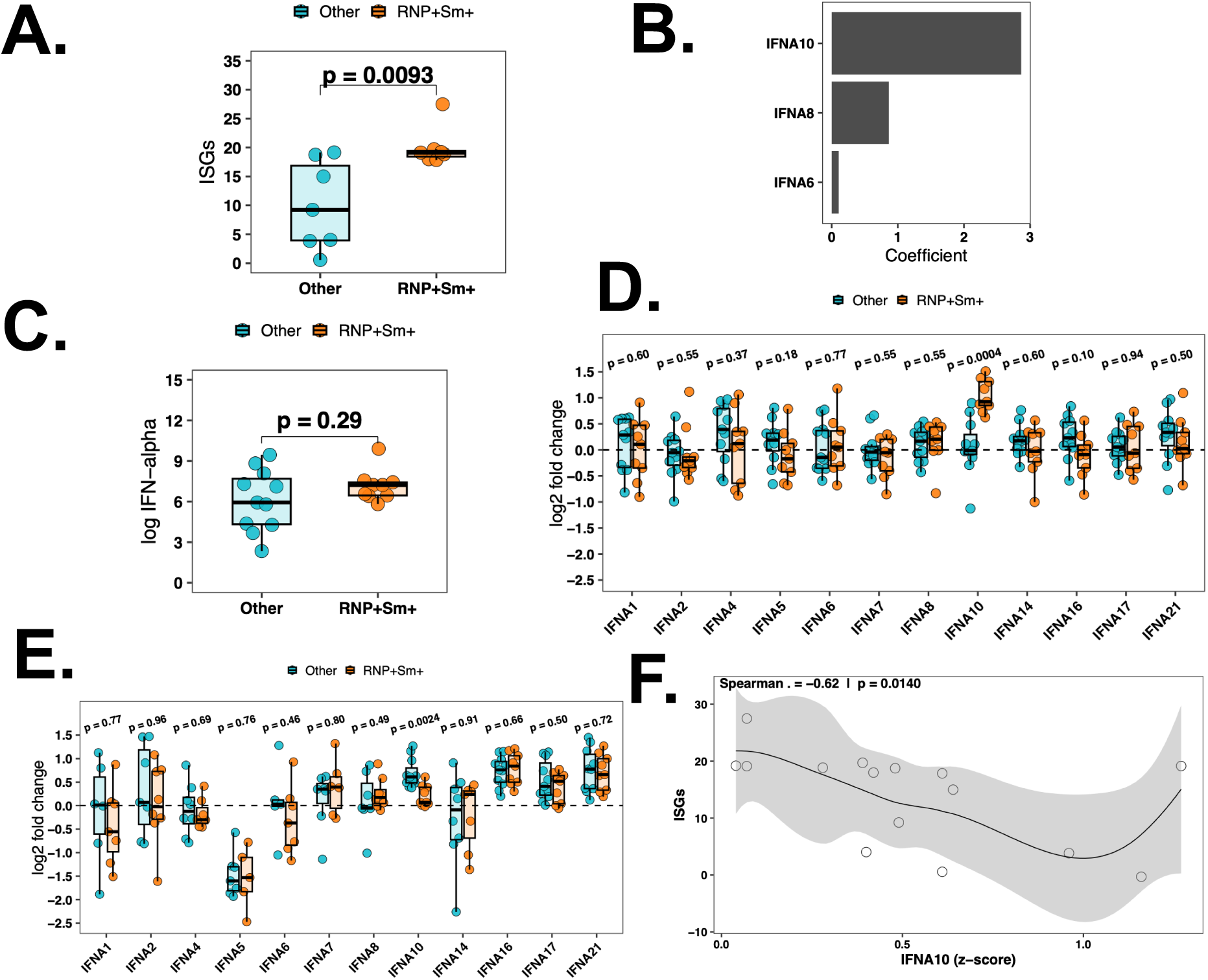
**A-F:** Elevated Interferon-α (IFNA)-10 and ISG signatures in RNP⁺Sm⁺ lupus. **A.** Interferon-stimulated gene (ISG) expression in RNP⁺Sm⁺ (n=9) versus other lupus subgroups (n=11). **B.** Top IFNA subunits most predictive of ISG expression, identified by penalised lasso regression analysis of transcriptomic data. **C.** Comparison of total serum IFNA concentrations between RNP⁺Sm⁺ and other lupus patients. **D.** Bar plots showing transcriptomic expression for individual IFNA subunit mRNA expression in the two groups, expressed as fold change relative to the control group. **E.** Bar plots depicting IgG autoreactivity against individual IFNA subunits, as measured by proteome microarray, in RNP⁺Sm⁺ lupus subjects compared to other lupus subjects. Data are shown as fold change relative to the control group. **F.** Spearman correlations illustrating relationships between IgG autoreactivity to IFNA subunits and ISG expression across the cohort. *Pairwise group p-values were calculated using linear regression adjusted for age, ethnicity, and baseline SLEDAI*.

Given previous reports that endogenous anti-IFNA autoantibodies are associated with reduced SLE disease severity (31), we next assessed IgG autoreactivity to IFN-α subtypes using proteome microarrays from the BEAT-Lupus cohort. RNP⁺Sm⁺ patients showed significantly lower autoreactivity to IFNA10 (p=0.0024; Figure 7E), and IFNA10-specific autoreactivity was inversely correlated with ISG expression (R = −0.62, p=0.0140; Figure 7F).

## Discussion

Our study identifies RNP⁺Sm⁺ co-positivity as a high-risk, mechanistically distinct, and clinically aggressive myeloid-dominant SLE endotype. This endotype represents more than one-third of treatment-refractory patients across three independent RCTs and an observational cohort. The RNP⁺Sm⁺ signature was validated across datasets with variable multi-omic depth, supporting its robustness as a serological marker of underlying biological heterogeneity. Three-way comparisons confirmed that single-positive (RNP or Sm alone) patients were phenotypically indistinguishable from double-negative controls across inflammatory markers, with cytokine levels demonstrating an ordered gradient from Neither→ Either⁺ → Both⁺, demonstrating that co-positivity—not single-antibody status—drives this distinct biology.

Patients within this endotype were more likely to be refractory to first-line therapy—a disparity most pronounced among Black patients, aligning with established associations between anti-RNP positivity and more severe lupus in this population (32). These patients subsequently exhibited poor clinical responses to second-line B-cell-depleting therapies, including rituximab and obinutuzumab. While alternative treatments such as belimumab and cyclophosphamide achieved response rates comparable to other serological groups, the failure of belimumab to yield superior outcomes in RNP⁺Sm⁺ patients is particularly striking given their elevated BAFF signature and myeloid-dominant inflammatory profile, suggesting that conventional B-cell-targeted and cytotoxic strategies are insufficient to address the underlying pathobiology of this endotype. This therapeutic resistance necessitated higher corticosteroid doses and resulted in significantly greater cumulative organ damage. Notably, Black patients with SLE—who comprised the majority of this endotype—have previously demonstrated reduced responses to B-cell-targeted therapies, including belimumab (33–35). Beyond treatment refractoriness, this endotype demonstrated specific propensity for severe phenotypic manifestations, including systemic vasculitis, lupus nephritis, and enteritis. Disease flares were consistently marked by pronounced hyperferritinaemia, establishing ferritin as a disease-specific biomarker that reflects the underlying myeloid-dominant inflammatory driver.

Despite achieving adequate peripheral CD20⁺ B-cell depletion, rituximab-treated patients failed to achieve sustained disease control, exhibiting accelerated B-cell repopulation driven by an early rebound in circulating BAFF alongside persistent elevation of pro-inflammatory cytokines. Critically, the absence of durable clinical response—even in the presence of successful B-cell clearance—coupled with rapid B-cell return and sustained myeloid cytokine production, suggests that B-cell-depletion strategies are insufficient for this endotype. These findings establish RNP⁺Sm⁺ status as a critical theragnostic biomarker that should steer researchers and clinicians away from reliance on B-cell-depletion therapies. Instead, this endotype necessitates alternative therapeutic paradigms directed at the interferon-dominated and myeloid-skewed inflammatory circuits that sustain disease activity. Potential strategies include interferon-pathway inhibitors, TLR7/8 or TLR4 antagonists, and the repurposing of downstream cytokine blockers such as IL-6 receptor antagonists.

Mechanistically, the RNP⁺Sm⁺ endotype reflects coordinated B-cell and myeloid activation driven by Sm/RNP immune complexes acting on the BCR-TLR7 axis within an interferon-dominated milieu (36, 37). Sm/RNP complexes are internalised via autoreactive BCRs and engage endosomal TLR7(36), triggering MyD88-dependent activation of IRF7 and NF-κB (38) and initiating type I IFN production (39, 40). While chronic type I IFN initially imposes BCR hyporesponsiveness on autoreactive B cells (41, 42), sustained TLR7 signalling coupled with cognate T-cell help via CD40L licenses full B-cell activation (36, 41), and promotes differentiation into T-bet⁺ CD11c⁺ DN2 memory B cells with glycolytic reprogramming (36, 37, 43), which we have found significantly enriched in RNP⁺Sm⁺ patients (44). DN2 cells preferentially differentiate into CD20^low^ plasmablasts and long-lived plasma cells (44), providing a direct mechanistic explanation for why CD20-directed B-cell depletion fails to yield durable clinical benefit in this endotype. Consistent with this framework, phase II data with the dual TLR7/8 antagonist enpatoran showed rapid suppression of interferon-stimulated gene expression within two weeks of treatment, supporting TLR7/8 inhibition as a rational strategy for RNP⁺Sm⁺ disease.

Beyond this initiating circuit, interconnected feed-forward loops amplify inflammation in the RNP⁺Sm⁺ endotype. Elevated IFN-γ levels and expanded IFN-γ-secreting T cells provide potent help for autoreactive B cells (45, 46) while the resulting immune complexes further augment type I IFN activity and BAFF secretion from myeloid and dendritic cells (47, 48). BAFF supports survival and expansion of DN2 cells, which express high levels of BAFF receptor (27, 49), explaining both the rapid B-cell repopulation after rituximab and the reduction in DN2 cells we observed with belimumab (27). Anti-Sm and anti-RNP immune complexes synergistically activate monocytes and macrophages to release IL-6 and TNF-α (50), providing additional help for B and T cells while driving tissue-directed inflammation. Our transcriptomic and proteomic data align with this model, revealing dominant monocyte and neutrophil signatures accompanied by elevated tissue-damaging mediators (TWEAK, S100A12, MMP-10) and myeloid-attracting chemokines (CCL2, CXCL10, CXCL9).

Consistent with with myeloid-driven pathology, pathway enrichment identified upregulation of TLR4-associated gene signatures, and flow cytometric validation confirmed increased TLR4 surface expression on intermediate monocytes during active flares (51). The marked elevation in serum ferritin observed during flares in RNP⁺Sm⁺ patients provides a readily accessible serological correlate of macrophage activation (52). Notably, despite profound hyperferritinaemia, only one patient fulfilled clinical criteria for haemophagocytic lymphohistiocytosis (HLH). Ferritin elevation, combined with normal or modestly elevated CRP, indicates that inflammatory burden in this endotype is mediated by type I/II interferons and myeloid cytokines rather than IL-1-driven acute phase pathways (53, 54). The strong correlation between ferritin and SLEDAI scores exclusively within the RNP⁺Sm⁺ group—not observed in other serological subgroups—establishes ferritin as a disease-specific activity marker for this endotype. These data support a model in which anti-RNP/Sm immune complexes amplify innate immunity through endosomal TLR7 or surface TLR4 signalling to sustain cytokine-driven pathology.

The RNP⁺Sm⁺ endotype exhibits qualitative differences in type I IFN signalling, with ISG induction most closely linked to IFNA10 mRNA rather than total circulating IFN-α. RNP⁺Sm⁺ patients showed selectively reduced anti-IFNA10 IgG autoreactivity that inversely correlated with ISG expression (31), suggesting loss of a physiological brake on IFNA10 activity. Complementing these immune signatures, functional pathway analyses indicated upregulation of tryptophan-arginine-proline metabolism and increased oxidative phosphorylation (55–60), consistent with IFN/TLR7-skewed immunity requiring enhanced bioenergetic and biosynthetic support for DN2/plasmablast differentiation and myeloid activation.

Anti-RNP and anti-Sm antibodies target distinct epitopes within the U1 small nuclear ribonucleoprotein complex: anti-Sm antibodies recognise core SmB and SmD proteins, whereas anti-RNP antibodies target U1-snRNP-associated proteins (U1-70kDa, A, and C) (61). Notably, co-positivity for both anti-RNP and anti-Sm was required to define this endotype, as single-positive patients displayed phenotypes indistinguishable from double-negative controls. This contrasts with previous reports associating single RNP or Sm positivity with B-cell hyperactivity characterised by elevated plasmablast-to-memory B-cell ratios and DN2 expansion (44, 62). HuProt proteome microarray profiling confirmed epitope spreading of IgG autoantibodies to other spliceosome and RNA processing proteins, with trend analyses demonstrating stepwise increases in autoreactivity paralleling cytokine escalation. Beyond spliceosomal targets, IgG autoreactivity extended to circadian-metabolic regulators (SIRT1, NCOA1, SREBF1, NR1D1/REV-ERBα, PER1). Anti-SIRT1 autoreactivity was particularly prominent and has not been previously described in SLE (63). Given that circadian dysregulation promotes cytokine production and renal injury in lupus (64, 65), this expanded autoreactivity may contribute to the heightened flare risk and nephritis susceptibility in RNP⁺Sm⁺ disease, though whether it causally drives pathology or reflects broader immune dysregulation requires investigation.

In summary, this study identifies RNP⁺Sm⁺ co-positivity as a prevalent, myeloid-driven inflammatory SLE endotype characterised by resistance to treatment including B cell depletion therapy. Mechanistically, this endotype is sustained by TLR7-driven DN2/plasmablast expansion, BAFF-dependent B-cell repopulation, and persistent inflammatory cytokine production, explaining poor rituximab responses despite adequate CD20⁺ depletion. Its disproportionate prevalence among Black patients—who experience higher disease severity and treatment refractoriness—underscores critical health disparities and unmet therapeutic need. In addition, this endotype is further defined by marked hyperferritinaemia, which correlates tightly and exclusively with disease activity in RNP⁺Sm⁺ patients, establishing ferritin as a readily measurable biomarker of flare and myeloid-driven inflammation in this subgroup. These findings establish anti-RNP/Sm co-positivity as an endotypic marker and ferritin as its companion disease activity biomarker to guide treatment selection using alternative strategies better aligned with the underlying interferon and myeloid-driven biology.

### Limitation of the Study

While the integration of data from multiple RCTs and observational cohorts confers robustness, certain limitations warrant consideration. First, the comparator (’Other’) group encompasses heterogeneous serological profiles, including isolated anti-dsDNA positivity, APS-predominant profiles, and seronegative patients. However, our BEST analysis systematically evaluated all possible antibody combinations—spanning anti-dsDNA, APS markers (IgG anti-cardiolipin, anti-β2-glycoprotein I), and conventional ENAs—and unequivocally identified RNP⁺Sm⁺ co-positivity as the subset with the strongest correlation to multivariate biological heterogeneity. Furthermore, comparative PERMANOVA modelling confirmed that alternative serological groupings based on anti-dsDNA or antiphospholipid antibodies explained significantly less variance in the global molecular landscape than the RNP⁺Sm⁺ signature.

Three-way comparisons demonstrated that single-positive patients were phenotypically indistinguishable from double-negative controls, and ordered gradient analyses confirmed stepwise increases in cytokine levels paralleling serological status, indicating that the observed differences are driven by dual positivity rather than artifacts of comparator heterogeneity. While conventional clinical assays used in these cohorts do not distinguish between individual RNP or Sm subunits, orthogonal HuProt proteomic analysis confirmed broad IgG enrichment against all major spliceosome components.

Second, high-dimensional data types, including proteomic profiling, cytokine measurements, and autoantigen arrays, were not uniformly available across all cohorts, potentially limiting the extrapolation of specific multi-omics signatures. Finally, although functional assays demonstrating enhanced TLR4 expression and inflammatory cytokine production in intermediate monocytes support a mechanistic basis for our findings, the cross-sectional design of the mechanistic studies precludes definitive causal inferences. Nevertheless, the convergence of serological, transcriptomic, proteomic, and clinical evidence from multiple independent sources provides compelling support for a biologically coherent myeloid-dominant SLE endotype defined by anti-RNP/Sm co-positivity.

## STAR★Methods

### Key resources table

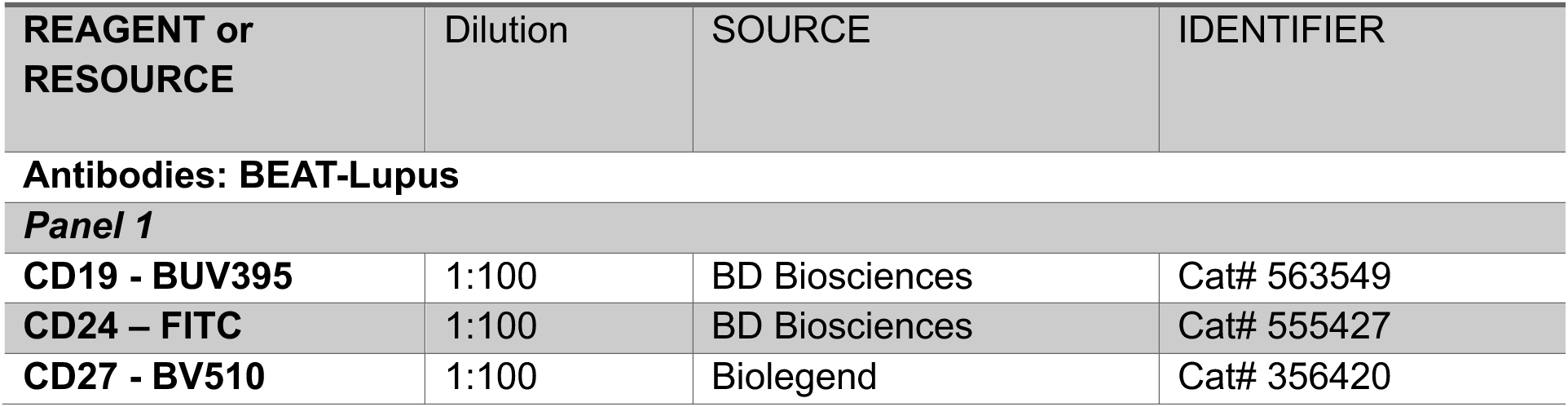
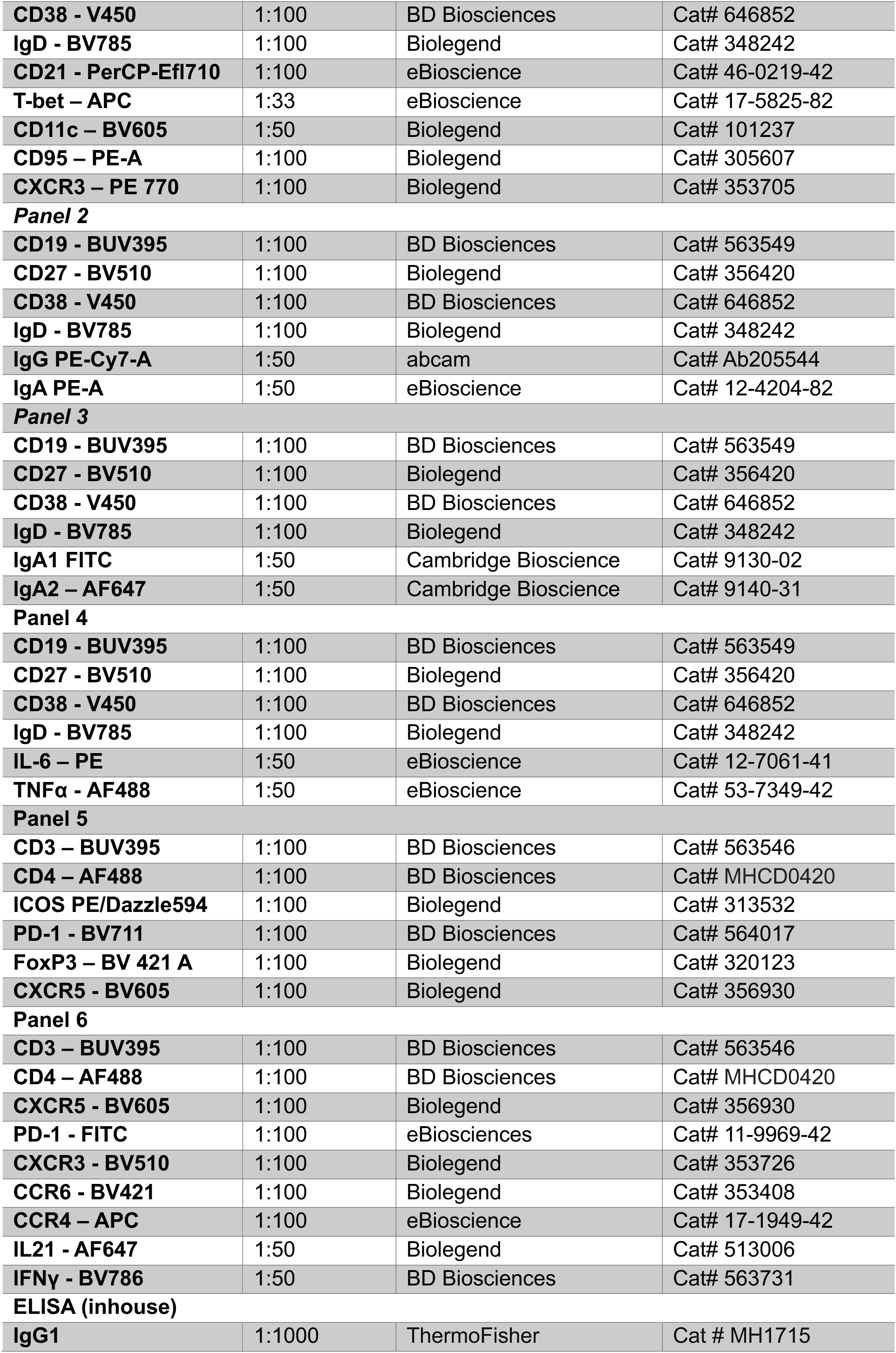
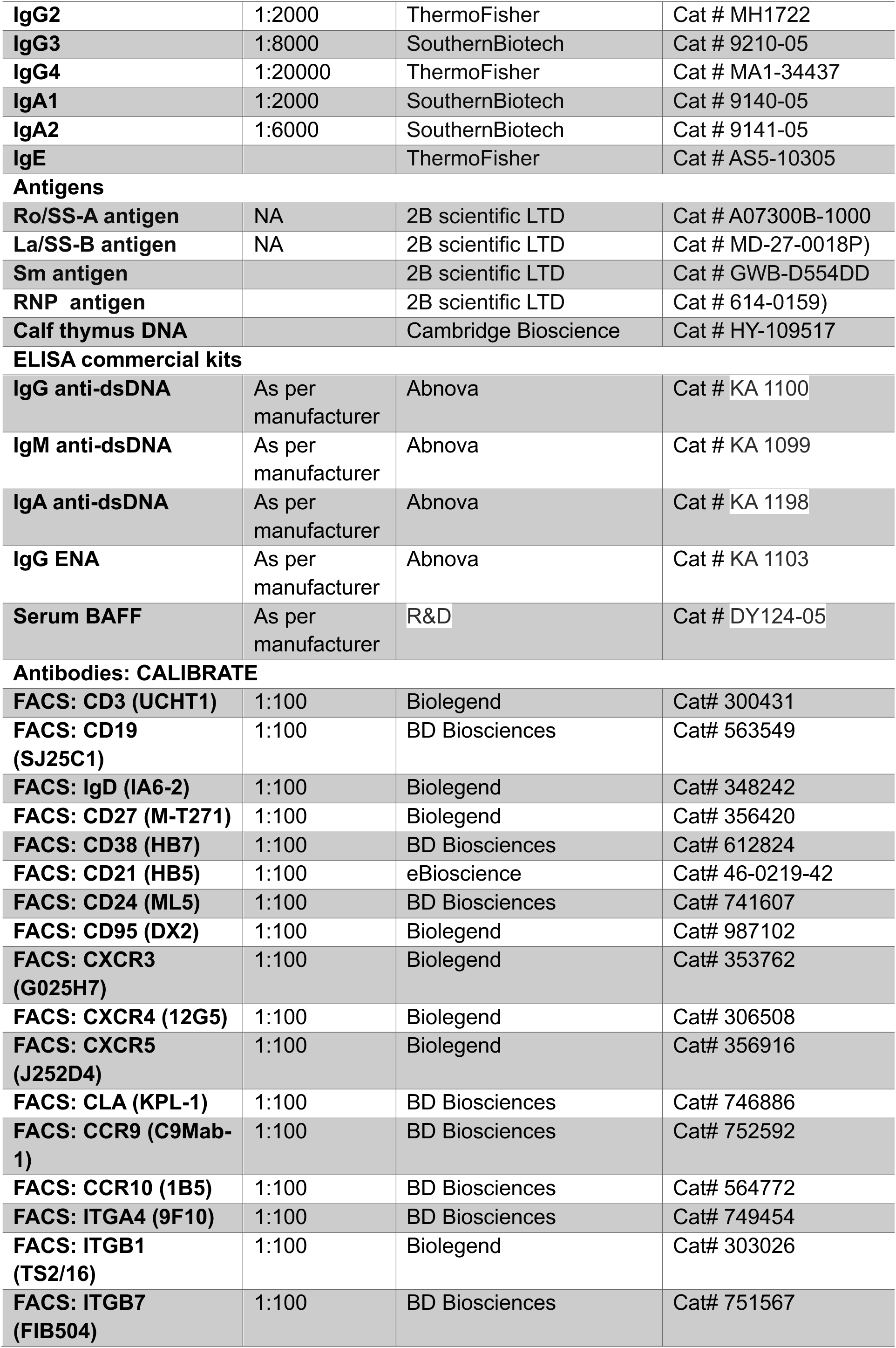
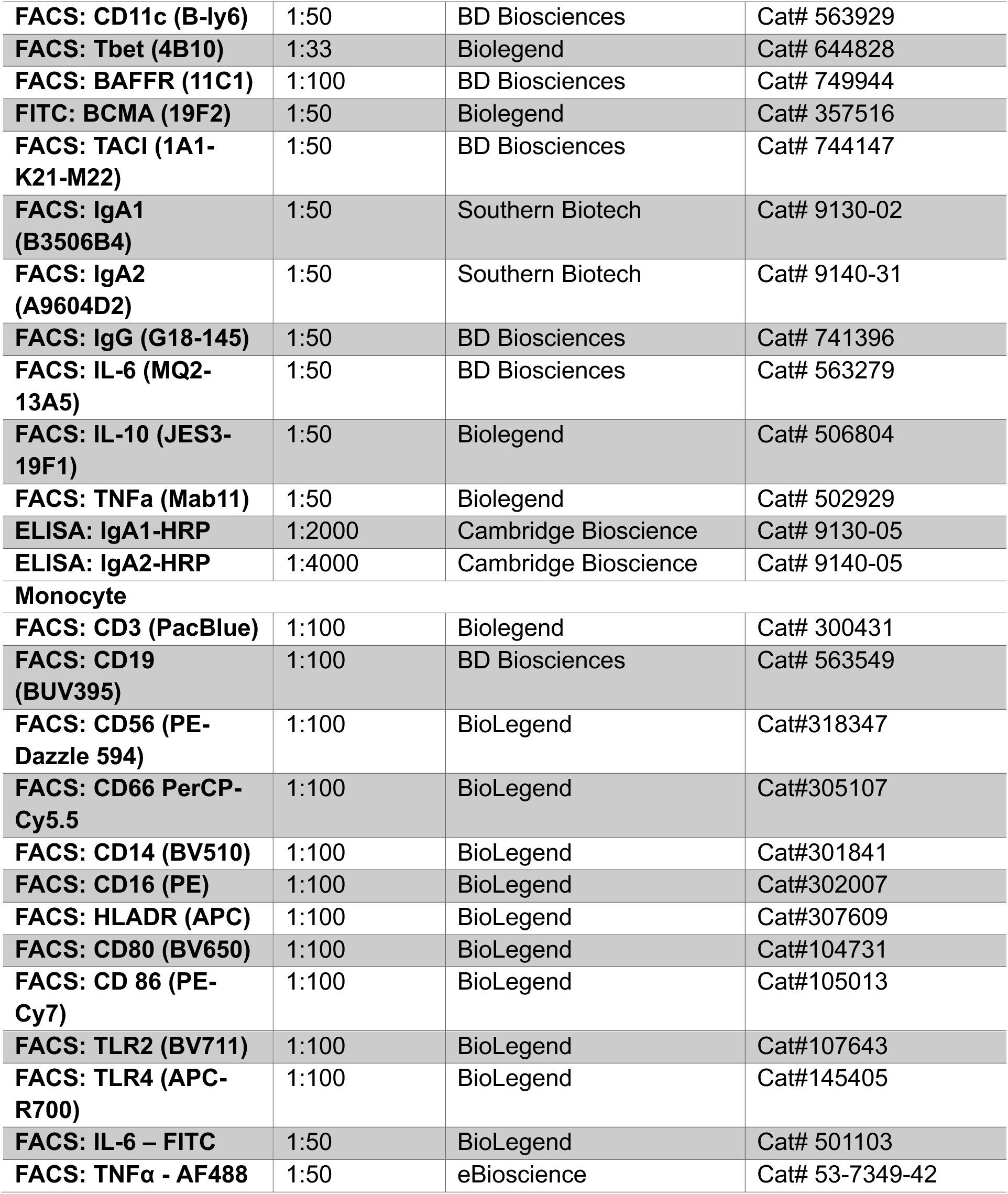

### Antibodies: BEAT-Lupus

#### Materials and Methods Study Cohorts

Various datasets were utilised in this analysis (Supplementary Dataset Section and Supplementary Table 1). The BEAT-LUPUS trial was a UK-based, multicentre, phase IIb, randomised, double-blind, placebo-controlled, parallel-group superiority trial with 52-week follow-up (12). Patients with SLE received B-cell depletion with rituximab, followed by randomisation to either belimumab or placebo, administered 4–8 weeks after the first rituximab infusion. We also included data and blood samples from CALIBRATE, an open-label, multicentre, randomised clinical trial conducted by the Immune Tolerance Network in the USA, enrolling patients with recurrent or refractory lupus nephritis (13). All participants received rituximab and cyclophosphamide two weeks apart before being randomised to either belimumab or continuation of standard therapy. The third RCT was the ACCESS trial, a 52-week multicentre study comparing abatacept versus placebo; both arms of the trial were given cyclophosphamide according to the Euro-Lupus protocol followed by azathioprine (25).

In addition, we analysed an internal observational cohort from the UCLH Lupus Centre comprising consecutive new patients seen between June 2019 and June 2024, as well as our previously published observational data (30) (Supplementary Table 1). External validation was performed using an independent cohort from Cambridge University Hospitals (CUH; November 2023 to November 2025). Finally, we interrogated publicly available NCBI GEO cohort GSE72326 with autoantibody metadata (66).

All contributing studies had prior research ethics approval.

### Definition of response and flares

#### Clinical Response

Two complementary response definitions were applied depending on cohort and analysis: Major Clinical Response (MCR) was defined as:

- Reduction of all BILAG-2004 Grade A or B domain scores to Grade C (stable inactive) or D (previously active but now inactive)
- Prednisolone dose ≤7.5 mg/day
- Modified SLEDAI-2K score ≤2 (excluding the anti-dsDNA antibody component) Persistent Low Disease Activity (LDA) required:
- No BILAG-2004 Grade A or B scores in any organ system at two consecutive visits (≥4 weeks apart)
- All organ systems scored as C, D, or E (never active)
- Prednisolone dose ≤5 mg/day (LDA5) or ≤7.5 mg/day (LDA7.5) at both visits
- Stable immunosuppressant and hydroxychloroquine doses for the preceding 12 weeks

Response timepoints varied by cohort:

- BEAT-Lupus trial: Week 52 post-randomisation (following rituximab at baseline)
- CALIBRATE trial: Week 48 post-randomisation (following rituximab and cyclophosphamide induction)
- ACCESS trial: Week 24 post-randomisation (following Euro-Lupus cyclophosphamide protocol)
- UCLH observational cohort: Week 24 for rituximab, belimumab, and obinutuzumab treatment cohorts

#### Flare Definition

Moderate-to-severe flares were defined as new or worsening disease activity requiring treatment escalation, defined as:

- BILAG-2004 Grade A (severe, active disease) in any organ system, **OR**
- Two or more BILAG-2004 Grade B scores (moderate disease activity) in different organ systems

Flares required a change in treatment (increased corticosteroids, addition or escalation of immunosuppressants, or initiation of biologic therapy), unless refused by the patient. Time-to-flare analyses censored patients at last follow-up or death. For recurrent-event analyses, each qualifying flare was recorded as a separate event, with patients remaining at risk for subsequent flares throughout follow-up.

#### B-Cell Repopulation

B-cell repopulation following rituximab or obinutuzumab was defined as CD19⁺ B-cell counts >10 cells/μL on two consecutive visits. Time-to-repopulation was calculated from the date of first B-cell-depleting agent administration to the first visit meeting this threshold.

## Method details

### Clinical and conventional laboratory data

Clinical and conventional laboratory data were abstracted from electronic health records or the relevant trial platforms – www.itntrialshare.org and NCBI.

### Antibodies and ELISA

Conventional autoantibody profiles were derived from the central laboratory data provided within the clinical trial platforms and standard diagnostic testing for the observational cohorts. Across these datasets, ‘anti-RNP’ and ‘anti-Sm’ denote reactivity determined by standard clinical solid-phase assays targeting the U1-snRNP complex and SmB/D proteins, respectively, without discrimination of individual subunit specificities. Other SLE-associated specificities, including anti-ribosomal P, anti-chromatin, and anti-C1q antibodies, are not routinely quantified in standard clinical practice nor reported across the trial datasets; consequently, these variables were excluded.

Serum antibody concentrations were quantified by ELISA. For IgA1, IgA2, IgG subclasses, and IgE, in-house ELISAs were used to detect anti-dsDNA and ENA antibodies according to previously published methods (26, 27). Data were reported as arbitrary units based on a standard curve generated from serially diluted control. Commercially available ELISA kits were employed to analyse IgG anti-dsDNA, ENA antibodies, and BAFF.

### Flow cytometry

Flow cytometric analysis was performed using both conventional (BEAT-Lupus) and spectral (CALIBRATE) platforms, following previously published established protocols for staining strategies and antibody selections (26, 27). For all samples, surface markers were stained for 30 minutes at 4°C, followed by overnight fixation and permeabilisation at 4°C and subsequent intracellular staining for 40 minutes at 4°C (67). Details of antibody clones, dilutions, and concentrations are listed in the Key Resources Table. Data acquisition was performed on a BD Fortessa for conventional flow cytometry and a Sony ID7000 for spectral flow. Data were pre-processed using FlowJo (v10), then exported to R for downstream analysis.

### Interferon signature and BAFF RNA expression

Whole blood (BEAT-Lupus) was collected into Tempus RNA tubes and stored at –80°C until processing. RNA was isolated using the Tempus RNA isolation kit, then reverse transcribed into cDNA. Quantitative real-time PCR was performed on an ABI Prism 7900HT, with gene expression normalized to GAPDH as a control. Seven interferon-stimulated genes (ISGs; ISG15, IFI44, RASD2, STAT1, SERPING1, BST2, and SP100) were measured, and each sample’s total interferon score was calculated as the summed standardized fold-change relative to the median of 20 healthy controls [Σ(RE_subject – RE_HC)/SD_HC, where RE is the relative expression]. Sub-signatures for interferon-α (ISG15, IFI44, RASD2) and interferon-β (STAT1, SERPING1, BST2, SP100) were also computed as previously described. BAFF gene expression in whole blood was quantified using the same qPCR protocol.

### Cytokines

For cytokines, we used Simoa Human Cytokine 6-Plex Panel-1 Advantage Kit (Quanterix, Billerica, MA, USA) to measure serum concentrations of IFN-γ, interleukin (IL)-10, IL-12p70, IL-17A, IL-6, and tumour necrosis factor (TNF), and to measure IFN-α we used the Simoa IFN-α Advantage Kit (100860, Quanterix, Billerica, MA, USA) on a Simoa HD-1 analyser.

### Inflammatory Proteomics

For inflammatory profiling, plasma samples were analysed with the Olink® Target 96 Inflammation Panel (Olink Proteomics, Uppsala, Sweden). This high-throughput assay simultaneously quantifies 92 inflammation-related proteins in each sample, utilising 1 μL of plasma. The platform leverages the proximity extension assay (PEA) technology: pairs of oligonucleotide-labelled antibodies specifically bind their target proteins, and only those binding in close proximity produce a unique reporter DNA sequence by DNA polymerization. These DNA barcodes are then amplified and quantified using high-throughput real-time PCR (Fluidigm Biomark HD system), providing normalised protein expression (NPX) values on a log2 scale. Internal controls and sample handling checks are incorporated throughout the protocol to ensure data quality and comparability across runs. Panel details, biomarker coverage, and additional quality information are available from the manufacturer @-https://olink.com/products/olink-target-96.

### HuProt™ protein microarrays

HuProt™ protein microarrays (v4.0; Cambridge Protein Arrays) were used to profile serum autoantibodies against >23,000 purified human proteins (>16,000 genes) printed in duplicate with batch QC by anti-GST staining; arrays were stored at −80 °C until use. Paired sera from BEAT-Lupus patients (screening and 12 months) plus a secondary-only negative-control slide were assayed for IgG and IgA. Arrays were blocked overnight at 4 °C in PBS/0.05% Tween-20/2% BSA, then incubated with sera diluted 1:1000 for 2 h at room temperature with gentle rocking, washed (3× PBS-Tween, then 2×5 min PBS-Tween), and probed for 2 h with Fc-specific anti-human IgG-546 (4 µg/mL), α-chain–specific anti-human IgA-488 (8 µg/mL) and anti-GST-650 (1 µg/mL), followed by identical washes, ddH₂O rinse and spin-dry. Slides were scanned on a Tecan LS400 (10 µm; 532 nm IgG, 488 nm IgA, 633 nm GST), features were quantified in GenePix 7, duplicate spots averaged, and sample signals background-corrected by subtracting the matched negative-control signal and log₂-transformed. For each protein, a z-score across the array distribution was computed to assess significance, and an Interaction Score was derived to account for protein content (i = [Prot s-c / GST sam] × 10). Data were filtered using prespecified QC thresholds (duplicate rsd < 0.35, signal-to-noise > 2.5, and sample/control ratio > 2), with artefact-flagged spots excluded; remaining proteins were ranked by z-score for downstream analysis and visualisation.

For functional and pathway enrichment, differential autoantigen reactivities were pre-ranked by z-score and analysed in R (v4.5.0) using the limma package (v3.58.0), with competitive gene set testing by CAMERA and gene set collections obtained from MSigDB (v2025.1.Hs, 2025) (Gene Ontology Biological Process, Gene Ontology Cellular Component, and Reactome pathways) (68–72).

### Bulk RNA-sequencing

Total RNA of BEAT-Lupus was extracted from whole blood collected in EDTA tubes using the RNeasy 96 kit (cat. no. 74192; Qiagen) with on-column DNase I digestion (RNase-Free DNase Set, cat. no. 79254; Qiagen) according to the manufacturer’s protocol. RNA concentration was measured by fluorometry, and integrity was assessed on an Agilent Bioanalyzer. Poly(A)-selected, strand-specific, dUTP-based (reverse-stranded) libraries were constructed, quality-checked for size distribution (peak ∼300–500 bp) and molarity, pooled equimolarly, and sequenced on an Illumina NovaSeq 6000 S4 flow cell (2×150 bp) to a median depth of ∼40–50 million paired-end reads per sample. Where applicable, globin-blocking oligonucleotides were included during library preparation to mitigate hemoglobin mRNA carryover from whole blood.

Sequencing quality control (QC) required RNA integrity RIN ≥ 7 (or DV200 ≥ 30% for partially degraded RNA), acceptable library size distribution, ≥85% bases with Q≥30, and minimal adapter contamination. Adapter and low-quality base trimming was performed with fastp (v0.23.2)(73). Reads were aligned to GRCh38 with STAR (v2.7.11a; two-pass mode) using GENCODE v45 gene models(74, 75). Alignment QC included total and unique mapping rates, rRNA content, insert size, and 5′–3′ coverage bias. Gene-level quantification used featureCounts (v2.1.1) with secondary/supplementary and multi-mapping reads excluded (76). Samples were further assessed for library size, % uniquely mapped reads, mitochondrial and rRNA content, duplication metrics, 5′–3′ bias, and by PCA on variance-stabilized counts to identify outliers and batch effects. Pre-defined exclusion criteria were overall mapping <50% and/or multi-metric outlier status.

Downstream analysis was performed in R (v4.5.0) using DESeq2 (v1.49.4) (77). Low-abundance genes were filtered a priori (e.g., counts ≥10 in ≥20% of samples) before modeling. Size-factor normalization, dispersion estimation, and Wald tests followed the DESeq2 workflow; the false discovery rate (FDR) was controlled at 5% (Benjamini–Hochberg). Known batch variables (e.g., sequencing lane, library-prep batch) were included as covariates in the design formula. For visualization only, removeBatchEffect (limma) was applied to transformed data. Differential expression was called at FDR < 0.05; genes with |log₂FC| > 0.5 were highlighted for pathway analyses.

Module-level expression (M-scores) for immune-related gene signatures were computed from bulk RNA-seq using a published approach (18). Genes were mapped to 606 pre-defined blood transcriptional modules from the combined Li et al. (BTM) and Chaussabel collections using the tmod (v0.48.2) R package (78, 79). For each individual, gene expression was standardized to z-scores against healthy control distributions, and module-level M-scores were calculated as the average z-score for all genes within each module.

Immune cell composition was inferred from bulk RNA-seq using CIBERSORTx, a widely validated digital cytometry framework that estimates the relative abundance of 22 immune cell subsets from whole-transcriptome profiles based on signature gene expression matrices (80).

Pathway and functional enrichment of differentially expressed genes relied on over-representation and pre-ranked gene set enrichment analyses using the latest clusterProfiler (v4.17.0) and ReactomePA (v1.53.0) (81, 82), mapping identifiers via org.Hs.eg.db (v3.20.0). Analyses referenced Reactome, KEGG, and MSigDB v2025.1.Hs collection (70, 83), applying FDR control throughout.

## Statistical analysis

All analyses were conducted in R (v4.5.0, macOS). Two-sided p values and 95%CIs are reported. Continuous variables were compared between groups with Student’s t test or Mann–Whitney U test; for non-parametric pairwise contrasts following a global test we used Dunn’s test with Benjamini–Hochberg adjustment. To account for confounding, multivariable linear regression was employed with covariate sets selected based on the specific analysis context. Phenotypic comparisons (cytokines, cell subsets, gene signatures) were adjusted for baseline SLEDAI-2K scores, age, and ethnicity to isolate endotype-specific biological associations independent of disease activity. For clinical outcome analyses (time-to-flare, treatment response), models were additionally adjusted for baseline medication use (prednisolone and immunosuppressant dosage) and the presence of lupus nephritis to control for prognostic factors affecting therapeutic efficacy.

Categorical outcomes used Fisher’s exact test and, where appropriate, logistic regression fit via glm (family = binomial, logit link) to estimate odds ratios (ORs) with 95%CIs. For models at risk of small-sample bias or (quasi-)complete separation—e.g., the prespecified treatment × endotype interaction—we used Bayesian logistic regression via bayesglm (arm v1.14.4) with default weakly informative priors, to stabilise coefficient estimates and ensure finite, conservative ORs compared with maximum-likelihood glm. Multivariable odds ratios for organ involvement were estimated by L1-penalised logistic regression using the glmnet package (v4.1-10), with the optimal shrinkage parameter (λ) chosen via cross-validation (84). Model coefficients are reported as odds ratios with 95%CIs. Associations were summarised by Spearman’s ρ with 95%CIs from bootstrapping. Unless stated otherwise, multiplicity was controlled within analysis families using Benjamini–Hochberg FDR (5%). Missing data were handled by complete-case analysis with multiple imputation (mice v3.18.0) as a sensitivity check; influential points were screened via leverage/Cook’s distance (85).

To identify the autoantibody signature most strongly aligned with global molecular heterogeneity, we integrated multimodal data using MOFA+ (MOFA2 v1.20.2) (28), deriving a low-dimensional latent representation of inter-patient variation. Input views comprised clinical characteristics and conventional laboratory parameters, cytokine profiles, transcriptomic data including interferon/ISG signatures, and T- and B-cell immunophenotyping. Variables were Z-score scaled within view prior to model fitting, and MOFA+ was fitted using Gaussian likelihoods. The model was initialised with K = 20 latent factors and trained using variational inference until convergence. We retained factors explaining ≥2% variance in at least one view (total n=9 factors); Factor scores were z-standardised and global inter-patient dissimilarity was computed as Euclidean distances using all retained MOFA factors.

The serological matrix comprised nine antibody specificities (Sm, RNP, Ro, La, IgG anti-cardiolipin, IgG anti-β2-glycoprotein I, IgG anti-dsDNA, and IgA1/IgA2 anti-dsDNA). To avoid unstable dissimilarities driven by very low-prevalence antibodies, we restricted the search space to antibody subsets in which each included specificity was present in at least 10% of patients (yielding 87 eligible subsets out of 511 possible; 2⁹−1). For each eligible subset, we computed a serology-based inter-patient dissimilarity matrix using binary Jaccard distance on positivity patterns (vegan::vegdist, method = “jaccard”, binary = TRUE). Subsets were ranked by the Spearman correlation (r) between the serology dissimilarity matrix and the global MOFA-derived dissimilarity matrix, and the subset maximizing r was selected (analogous to the BIOENV/BEST procedure; vegan v2.7-2) (29, 86). statistical significance of the correlation for the selected subset was assessed using a Mantel test with 9,999 permutations (vegan::mantel). As a sensitivity analysis, we repeated the MOFA–BEST pipeline using a stricter 20% prevalence threshold (≥10/52) for antibody inclusion, which yielded consistent identification of the RNP⁺Sm⁺ endotype. Further sensitivity analysis included, running MOFA–BEST pipeline after excluding the non-conventional IgA1/IgA2 anti-dsDNA antibodies from the serological panel. For external validation, we applied the same MOFA–BEST framework to three independent cohorts (CALIBRATE, ACCESS, and UCLH), adapting the input views to the omics and clinical data available in each dataset. Multivariate differences between resulting endotype groups were assessed using PERMANOVA (adonis2, vegan) with 9,999 permutations on the MOFA factor scores. As a sensitivity analysis, we additionally fitted PERMANOVA models for a set of predefined serological endotypes (IgG anti-dsDNA positivity alone; IgG anti-dsDNA positivity combined with any of RNP, Ro, La, Sm, IgG anti-cardiolipin, or IgG anti-β2-glycoprotein I), which showed weaker associations with the MOFA factor space than the data-driven RNP⁺Sm⁺ endotype.

Discriminative features were selected using ensemble: random forest with Boruta feature selection (87) (Boruta v9.0.0 (88) and cross-validated with penalised sparse partial least squares (sPLS)(89) by using the mixOmics package in R-v6.32.0 (90). Variables were ranked by their mean SHapley Additive exPlanations (SHAP) values(91).

To test for monotonic trends across ordered serology strata (Both⁺ → Either⁺ → Neither), we used the Jonckheere–Terpstra test, a non-parametric method for detecting directional alternatives across k groups. This was implemented using the coin package (v1.5-0; jonckheereTest) (92).

Time-to-event endpoints were analysed using Kaplan–Meier estimates (log-rank tests) and Cox proportional-hazards models with the survival (v3.8-4). Kaplan–Meier figures were rendered with survminer (v0.5.1). Proportional-hazards assumptions were evaluated via Schoenfeld residuals using cox.zph. For recurrent severe flares, we plotted mean cumulative function (MCF) curves using reda (v0.5.6) with 95%CIs, and estimated the hazard for the next flare using a Prentice–Williams–Peterson (PWP) total-time Cox model, stratified by event order and with patient-level robust (clustered) standard errors, adjusted for prespecified covariates (age, hydroxychloroquine, immunosupressants, and prednisolone dose) (93). Flare incidence rates were modelled with Poisson regression (stats::glm) reporting incidence rate ratios (IRRs); over-dispersion was assessed and a negative-binomial model (MASS v7.3-65; glm.nb) was substituted when indicated.

Treatment effect estimates were synthesised using random-effects meta-analysis with restricted maximum likelihood (REML) and the Hartung–Knapp–Sidik–Jonkman adjustment for confidence intervals, implemented in R v4.5.0 with metafor (v4.8-0; rma.uni, test = “knha”). Between-study heterogeneity was summarised by I² and τ² and displayed as forest plots. In sensitivity analyses, we also fit (i) a Bayesian normal–normal random-effects model (flat prior on the overall effect; half-Cauchy prior on the between-study SD) and (ii) a beta–binomial model with Jeffreys prior to stabilise estimates when responder counts were sparse; posterior medians and 95% credible intervals are reported for these analyses.

Longitudinal changes were analysed with linear mixed-effects models on the arcsine–square-root–transformed proportion, with endotype, visit (categorical), and their interaction as fixed effects and subject-level random intercepts (lme4 v1.1-37). Marginal means were back-transformed to the probability scale and compared with Tukey-adjusted contrasts using emmeans (v1.11.2-8). Truly binary repeated outcomes (present/absent) were, where applicable, modelled with mixed-effects logistic regression (glmer) with the same fixed-effect structure. Longitudinal continuous immunological measures (e.g. CD19⁺ B-cells, BAFF) were log-transformed and analysed with mixed-effects linear models including group, time, and group×time, with patient-level random intercepts; estimates are shown as geometric means with 95%CIs. When exactly two time points were compared, we used baseline-adjusted ANCOVA.

## Supplementary Materials

Index:

Supplementary Dataset Selection 2

**Supplementary Table:**

Supplementary Table 1 4

Supplementary Table 2 6

**Supplementary Figures**

Supplementary Figure 1 A-D: Immune clustering of SLE patients based on autoantibody profiles. 7

Supplementary Figure 2A-E: Serological and clinical characterisation of the RNP⁺Sm⁺ endotype across 4 cohorts 8

Supplementary Figure 3A-K: Immune clustering of lupus patients by anti-RNP and anti-Sm antibody status (RNP⁺Sm⁺) in BEAT-Lupus Trial Cohort at baseline.10

Supplementary Figure 4. Immune clustering of lupus patients by anti-RNP and anti-Sm antibody status (RNP⁺Sm⁺, single-positive, or double-negative)11

Supplementary Figure 5 A-D: Immune clustering of lupus patients by anti-RNP and anti-Sm antibody status (RNP⁺Sm⁺) in CALIBRATE Trial Validation Cohort at baseline.12

Supplementary Figure 6: Proportion of patients with various clinical manifestations in lupus patients with anti-RNP and anti-Sm antibodies (RNP⁺Sm⁺) compared with other lupus patients13

Supplementary Figure 7 A-D: Conventional laboratory markers and macrophage activation signatures at flare (UCLH and CUH cohorts)14

Supplementary Figure 8A–B: Proportion of patients achieving low disease activity on A. ≤5 mg/day and B. ≤7.5 mg/day prednisolone (LDA5) stratified by RNP⁺Sm⁺ endotype (positive for both anti-RNP and anti-Sm antibodies) versus all other patients in the Belimumab arm (n =26) of BEAT-Lupus trial.16

Supplementary Figure 9 A–C: RNP⁺Sm⁺ (positive for both anti-RNP and anti-Sm 17 antibodies) endotype does not differentiate response to belimumab or cyclophosphamide.

Supplementary Figure 10 A–E: Changes in CD19⁺ B-cell counts and serum BAFF (B-cell activating factor) stratified by endotype (RNP⁺Sm⁺: positive for both anti-RNP and anti-Sm) versus all other patients.19

Supplementary Figure 11: Changes in interferon-stimulated gene (ISG) expression 20 and baseline-elevated cytokines in the Beat-Lupus trial, stratified by autoantibody subgroup and treatment arm.

Supplementary Figure 12: Gating strategies to define intermediate, non-classical and classical monocytes.21

Supplementary Figure 13A–C: IgG (but not IgA)-associated autoreactivity predominates in spliceosome-related modules in RNP⁺Sm⁺ patients (BEAT-Lupus cohort).22

Supplementary Figure 14A–B: Jonckheere–Terpstra (JT) trend analysis of A. IgG- and B. IgA-associated spliceosome modules across ordered serology groups23

Supplementary Figure 15 A–F: Jonckheere–Terpstra (JT) trend analysis of 6 key cytokines of those patients with spliceosome modules available across ordered serology groups 24

Reference

## Acknowledgements

MRAS and MRE is partly funded through a block grant from the National Institute of Health Research to University College London Hospitals Biomedical Research Centre (BRC).

The authors gratefully acknowledge the CALIBRATE, BEAT-lupus, and ACCESS study teams. We are indebted to the patient participants in the CALIBRATE and BEAT-LUPUS trials, and other patients who contributed samples.

We thank Ellie Hawkins for recruiting patients and collecting the samples at University College London Hospitals NHS Foundation, Jamie Evans (the Rayne Flow Cytometry Facility, UCL) for flow cytometry technical support. We thank Dr Thomas McDonnell for advice about ELISA design and Mariea Parvaz, and Dr Liliana Santos for performing the flow cytometry and organising Huprot on the samples from the BEAT-lupus trial.

We also thank Prof. Coziana Ciurtin, Prof. Maria Leandro, and Ms. Rachel Allen, for providing the list of patients received obinutuzumab.

## Author Contributions

MRAS and MRE: Conceptualisation, methodology, formal analysis, data curation, visualisation, writing—original draft, writing—review and editing. CB and DM: Data curation, laboratory analysis, data access and analysis, writing—review and editing. DG: Conceptualisation, writing—review and editing. CC and ML: Resources (adolescent cohort), writing—review and editing. LC: Resources (CALIBRATE data), writing—review and editing. All authors approved the final manuscript.

## Resource Availability Statement

Data and codes are available upon reasonable request to the corresponding author, subject to approval by all co-authors and completion of a data access agreement.

