## Supplementary Materials for "Elevated ferritin as a companion biomarker of myeloid-driven inflammation in RNP/Sm co-positive, treatment-resistant SLE endotype"

### Supplementary Dataset Section:

**BEAT-Lupus (active SLE refractory to first-line therapy, requiring rituximab at the physician's discretion)** (1)

BEAT-Lupus was used as the primary discovery cohort, as it provided the most comprehensive and enriched set of variables, including clinical characteristics, conventional laboratory parameters, cytokine profiles, transcriptomics, serum interferon/ISG signatures, and T- and B-cell immunophenotyping. Longitudinal follow-up from BEAT-Lupus was also used to examine temporal changes in selected markers.

**CALIBRATE trial (refractory lupus nephritis)** (2)

CALIBRATE was included predominantly as a validation cohort. Full clinical data and conventional laboratory parameters were available, together with selected cytokine and flow-cytometry measurements based on shared sample availability. Because biomarker coverage was more limited than in BEAT-Lupus, CALIBRATE was not used for discovery clustering. Instead, it was employed to assess treatment effects and to validate key findings in an independent trial cohort predominantly comprising patients with refractory lupus nephritis, thereby allowing evaluation of the RNP$^+$Sm$^+$ endotype in a clinically distinct, renal-enriched population.

**ACCESS trial (lupus nephritis)** (3)

The ACCESS trial was included to evaluate treatment effects, particularly responses to cyclophosphamide-based regimens in a randomised controlled trial setting. Full clinical data and conventional laboratory parameters were available. This cohort was not used for clustering and was instead reserved for analyses of treatment response.

**UCLH internal cohort (validation)**

The UCLH (University College London Hospital) internal cohort was used primarily for external validation and comprised several subsets tailored to specific analyses; sample sizes therefore vary and are detailed in the relevant sections. For validation of clinical and conventional laboratory findings, we included patients first seen as new referrals to the UCLH lupus clinic between March 2019 and June 2025, a period chosen because implementation of the EPIC electronic health record enabled more precise and granular data capture.

For evaluation of treatment efficacy, we included patients who received rituximab, belimumab, or obinutuzumab as inpatients or in the dedicated infusion clinic between 2019 and 2025. These therapies were administered for a range of organ manifestations, but all patients had active disease, in line with national practice (BILAG A or two B scores, or SLEDAI >6).

For a subset of patients with available biobanked samples, proteomic and monocyte data were also incorporated to support mechanistic and biomarker analyses.

To estimate the incidence of refractory disease across $RNP^{+}Sm^{+}$ and others, we used the full UCLH SLE cohort with RNP/Sm results recorded in the electronic record.

**CUH cohort (external validation)**

The CUH (Cambridge University Hospital) cohort was included as an additional, geographically distinct external validation dataset to test the generalisability of refractory disease incidence across $RNP^{+}Sm^{+}$ and comparator groups in a different patient population. Data were extracted from November 2023 to November 2025, corresponding to the period of systematic data capture within the investigator’s CUH SLE cohort. Where available, ferritin measurements (not routinely performed at the same frequency as at UCLH) were used to provide supportive validation of ferritin as a biomarker of acute flare.

### Supplementary Table:

**Supplementary Table 1: List of datasets**

| Data set | Used for analysis | Sample Size |
|---|---|---|
| **BEAT-Lupus Trial**<br>**52 weeks, multicentred randomised controlled trial (RCT) by our group – comparing Belimumab versus placebo both after rituximab** | - Clinical data<br>- Conventional laboratories<br>- Various sub-and Isotypes of antibodies<br>- Interferon (IFN) signature genes and total scores<br>- Cytokines: BAFF, IL-6, IL-10, IL-12, IL-17, TNF-$\alpha$, IFN--$\alpha$, IFN-$\gamma$.<br>- RNA bulk sequencing<br>- Flowcytometry<br>- Huprot autoantigen repertoire | Clinical data and conventional lab –<br>Baseline (Belimumab arm, n = 26, Placebo arm, n = 26)<br>IFN, antibodies, and cytokines –<br>Baseline (Belimumab arm, n = 23, Placebo arm, n = 18),<br>Transcriptomics–<br>Baseline (Belimumab arm, n = 16, Placebo arm, n = 17)<br>Flowcytometry –<br>Baseline (Belimumab arm, n = 16, Placebo arm, n = 18)<br>Huprot microarray–<br>Baseline (Belimumab arm, n = 10, Placebo arm, n = 10)<br><br>*(Numbers may vary as some missing values and shown in respective result section)* |
| **CALIBRATE-trial**<br>**RCT done at USA– comparing Belimumab (RCB) plus standard of care versus standard of care alone (RC) both after rituximab and cyclophosphamide** | - Clinical data<br>- Conventional laboratories<br>- Antibodies (IgA1 and IgA2 anti-dsDNA)<br>- Cytokines: IL-6<br>- RNA bulk sequencing<br>- Flowcytometry | Clinical data and conventional lab –<br>Baseline (RCB arm, n = 21, Placebo arm, n = 22)<br>IFN, antibodies, and cytokines –<br>Baseline RCB arm, n = 16, RC arm, n = 16<br>Flowcytometry –<br>Baseline (RCB arm, n = 14, RC arm, n = 15) |
| **UCLH internal cohort** | Organ involvement and clinical data | Organ involvement<br>$RNP^{+}Sm^{+}$, n = 69; Other, n = 122 |

| | | |
|---|---|---|
| ***Internal cohort comprised new patients seen at the lupus clinic between June 2019 and June 2024*** | Treatment<br>Proteomics | <u>Treated with Rituximab</u><br>N = 62<br>RNP$^+$Sm$^+$, n = 27; Other, n = 35)<br><u>Treated with Belimumab</u><br>N = 42<br>RNP$^+$Sm$^+$, n = 18; Other, n = 24)<br><u>Treated with Obinutuzumab</u><br>N = 15<br>RNP$^+$Sm$^+$, n = 8; Other, n = 7)<br><u>Proteomes:</u><br>RNP$^+$Sm$^+$ n = 15, Other n = 15<br><u>Monocytes:</u><br>RNP$^+$Sm$^+$: active, n = 8, inactive n = 9, Other: active, n = 9, inactive n = 11.<br>Healthy control, n = 8 |
| **Previously observational published data from lupus unit of UCLH** | Flare to Rituximab | RNP$^+$Sm$^+$, n = 12; Other, n = 20 |
| **ACCESS-trial<br>52 weeks, multicentred RCT comparing Abatacept versus placebo both after cyclophosphamide followed by azathioprine** | Clinical, Transcriptomic | RNP$^+$Sm$^+$, n = 52; Other, n = 81 |
| **CUH investigator's lupus cohort between Nov 2023 and Nov 2025** | Clinical data | RNP$^+$Sm$^+$, n = 19; Other, n = 64 |
| **Publicly available NCBI-stored data (with metadata) GSE72326** | Transcriptomic | RNP$^+$Sm$^+$, n = 30; Other, n = 127 |

**Supplementary Table 2: PERMANOVA $R^2$ association with MOFA-derived molecular heterogeneity of the IgG autoantibodies combinations listed.**

| Serological grouping | BEAT-Lupus $R^2$ | CALIBRATE $R^2$ | ACCESS $R^2$ | UCLH $R^2$ |
|---|---|---|---|---|
| **$RNP^+Sm^+$** | 0.31 | 0.25 | 0.28 | 0.29 |
| **$dsDNA^+$ alone** | 0.07 | 0.11 | 0.12 | 0.04 |
| **$dsDNA^+RNP^+$** | 0.14 | 0.09 | 0.11 | 0.04 |
| **$dsDNA^+Ro^+$** | 0.12 | 0.16 | 0.14 | 0.22 |
| **$dsDNA^+La^+$** | 0.09 | 0.11 | 0.08 | 0.01 |
| **$dsDNA^+Sm^+$** | 0.15 | 0.05 | 0.02 | 0.09 |
| **$dsDNA^+aCL^+$** | 0.08 | 0.15 | 0.18 | 0.08 |
| **$dsDNA^+aB2GP^+$** | 0.11 | 0.04 | 0.10 | 0.05 |

### Supplementary Figures:

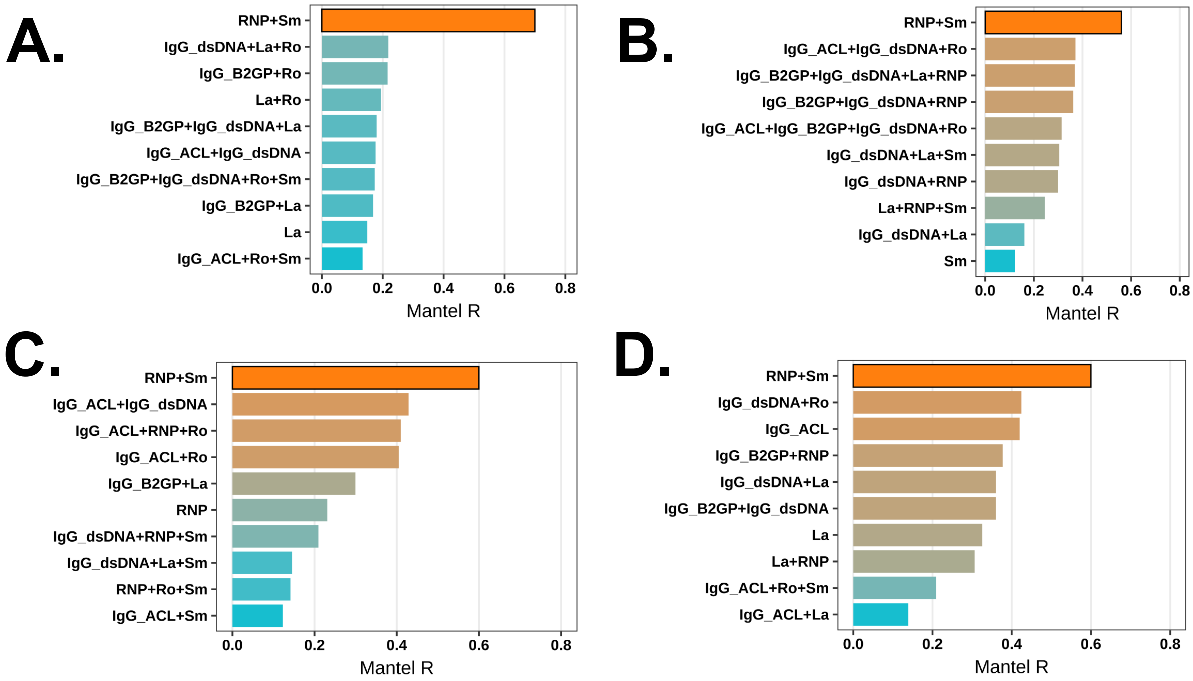


**Supplementary Figure 1 A–D: Immune clustering of SLE patients based on routinely available autoantibody profiles.** Multi-Omics Factor Analysis (MOFA) – Best Subset (BEST) serological analysis identifying autoantibody combinations that best capture global biological heterogeneity in four independent cohorts: **A.** BEAT-Lupus trial, **B.** CALIBRATE trial, **C.** ACCESS trial, and **D.** UCLH observational cohort. Bars show the Mantel correlation (Spearman method) between serology-based dissimilarity (Jaccard distance on antibody positivity patterns for seven specificities: Sm, RNP, Ro, La, IgG anti-cardiolipin, IgG anti-β2-glycoprotein I, and IgG anti-dsDNA) and the global MOFA-derived dissimilarity (Euclidean distance across latent factors). In the BEAT-Lupus discovery cohort (**A**), MOFA integrated clinical characteristics, laboratory parameters, cytokines, transcriptomics, ISG signatures, and deep immunophenotyping; replication cohorts (**B–D**) utilised variable-depth omics and clinical data (see Supplementary Table 1).

[$] *Cohort details: BEAT-Lupus (RNP$^+$Sm$^+$, n=17; Other, n=35); CALIBRATE (RNP$^+$Sm$^+$, n=12; Other, n=31); ACCESS (RNP$^+$Sm$^+$, n=52; Other, n=81); UCLH (RNP$^+$Sm$^+$, n=69; Other, n=122).*

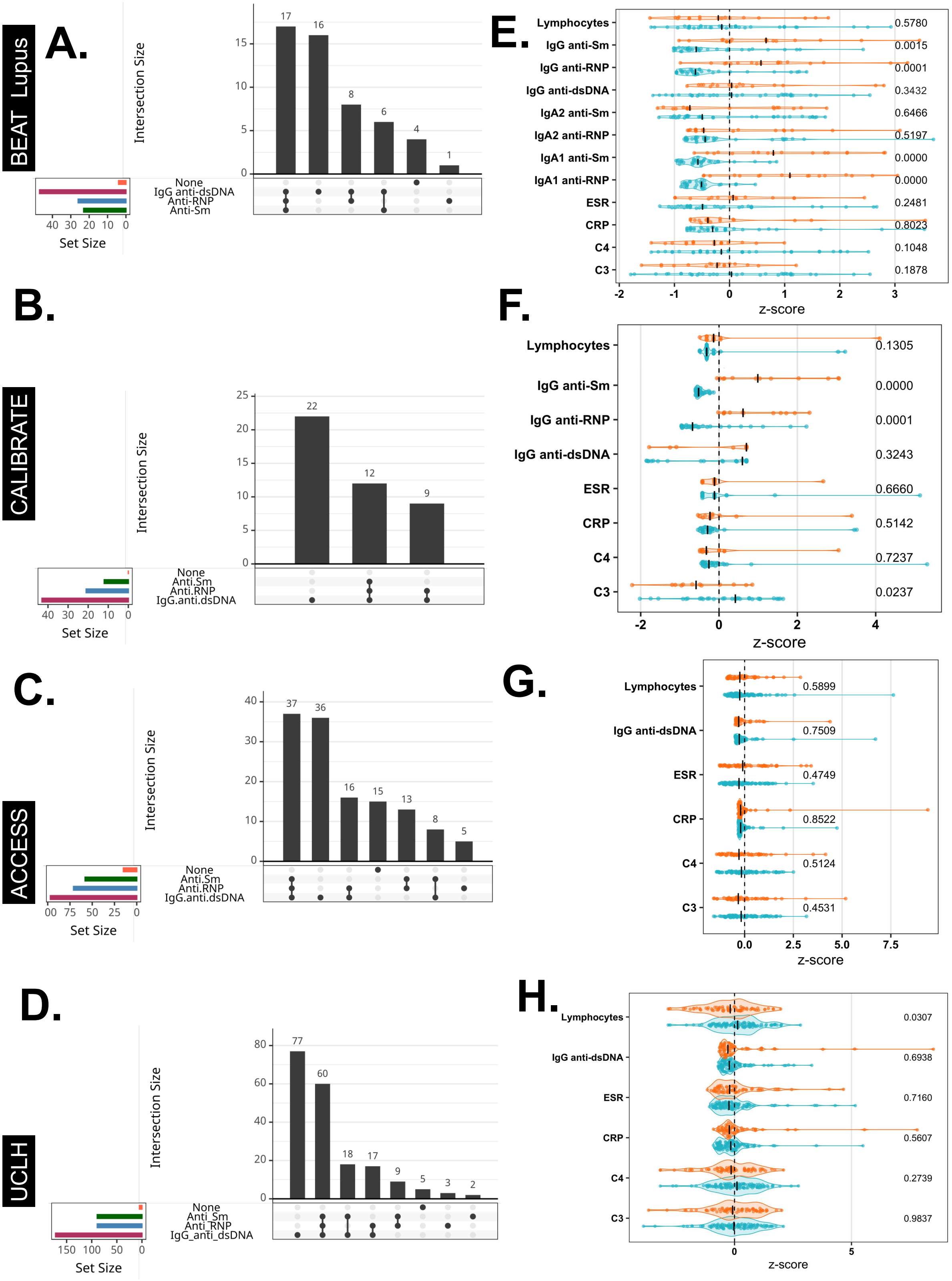


**Supplementary Figure 2A-H: Serological and clinical characterisation of the $RNP^{+}Sm^{+}$ endotype across 4 cohorts**[$]. UpSet plot displaying the intersection and frequency of autoantibody positivities, including IgG anti-dsDNA, anti-RNP, and anti-Sm at baseline **A.** BEAT-lupus, **A.** BEAT-Lupus trial, **B.** CALIBRATE trial, **C.** ACCESS trial, and **D.** UCLH.

Pairwise comparisons of conventional autoantibodies and disease activity-related laboratory markers (normalised as z-scores) between $RNP^+Sm^+$ and 'Other' patients across these 4 cohort **E-H**. p-values was assessed using the Mann–Whitney U test.

[$] *Clinical Trials - BEAT-Lupus trial ($RNP^+Sm^+$, n = 17; Other, n = 35), CALIBRATE trial ($RNP^+Sm^+$, n = 12; Other, n = 31), and ACCESS trial ($RNP^+Sm^+$, n = 52; Other, n = 81); Observational Internal cohort comprised new patients seen at the lupus clinic between June 2019 and June 2024 ($RNP^+Sm^+$, n = 69; Other, n = 122).*

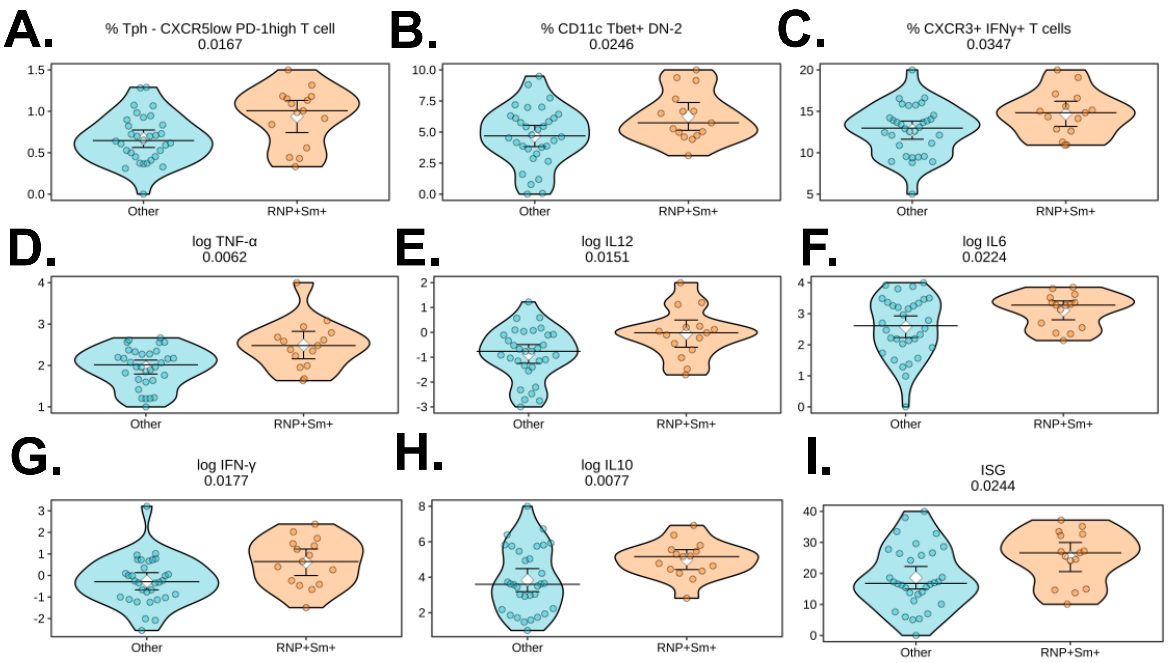


**Supplementary Figure 3A-I: Immune clustering of lupus patients by anti-RNP and anti-Sm antibody status ($RNP^+Sm^+$) in BEAT-Lupus Trial Cohort** (*n* = 52: $RNP^+Sm^+$, *n* = 17; Other, *n* = 35) **at baseline. A-I.** Pairwise comparisons of selected clinical and immunological variables between $RNP^+Sm^+$ patients and the remaining patients ('Other'). *p*-values for continuous variables were calculated using a linear regression model adjusted for disease activity, age, and ethnicity-Black vs. non-Black (unless otherwise indicated). Black horizontal lines represent the median, white diamonds indicate the mean, and vertical lines denote 95% confidence intervals. Cytokines were quantified by Simoa multiplex immunoassay; concentrations are reported in pg/mL and plotted as log10(pg/mL).

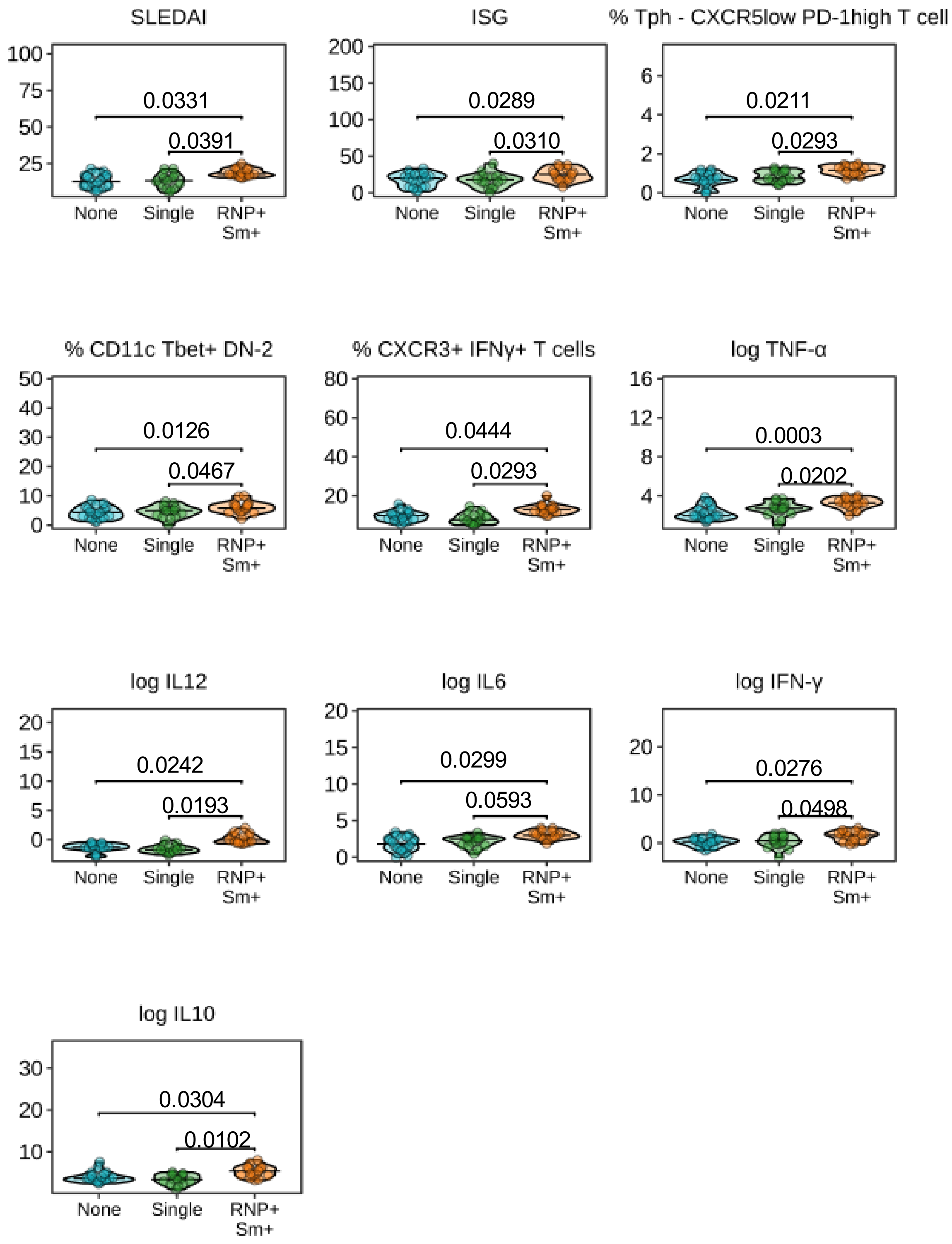


**Supplementary Figure 4. Immune clustering of lupus patients by anti-RNP and anti-Sm antibody status (RNP$^+$Sm$^+$, single-positive, or double-negative).** BEAT-Lupus trial cohort at baseline (n = 52; RNP$^+$Sm$^+$ n = 17, Other n = 35). Pairwise comparisons of selected clinical and immunological variables are shown. p-values for continuous variables were calculated using the Mann–Whitney U test. Cytokines were quantified by Simoa multiplex immunoassay; concentrations are reported in pg/mL and plotted as log10(pg/mL).

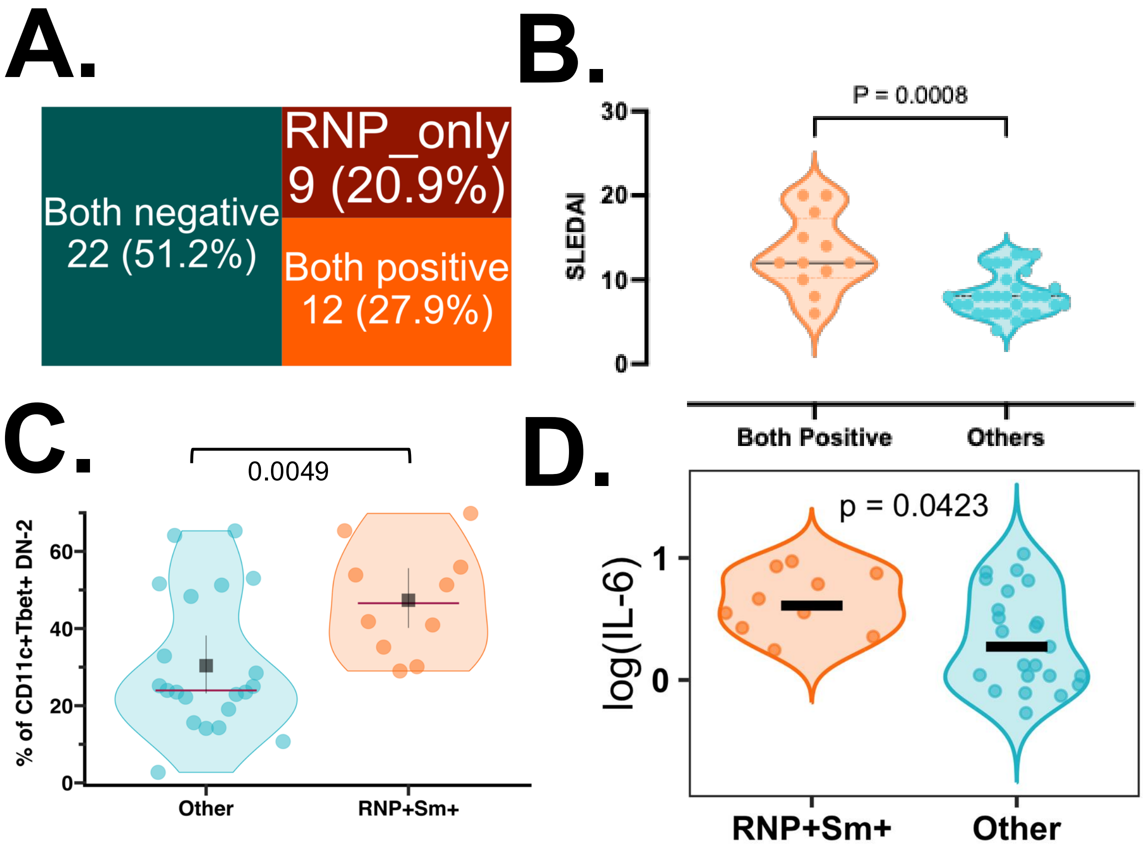


**Supplementary Figure 5 A-D: Immune clustering of lupus patients by anti-RNP and anti-Sm antibody status (RNP$^+$Sm$^+$) in CALIBRATE Trial Validation Cohort** (*n* = 43: RNP$^+$Sm$^+$, *n* = 12; Other, *n* = 31) **at baseline. A.** Stratification of patients by RNP$^+$Sm$^+$ status and other antibody profiles at baseline. **B.** Comparison of SLEDAI scores between RNP$^+$Sm$^+$ and other patients. **C.** Comparison of the proportion of CD11c$^+$T-bet$^+$ double-negative type 2 (DN2) B cells. **D.** Comparison of serum IL-6 levels. *p*-values were calculated using a linear regression model adjusted for age and ethnicity (SLEDAI was not adjusted for itself in panel B).

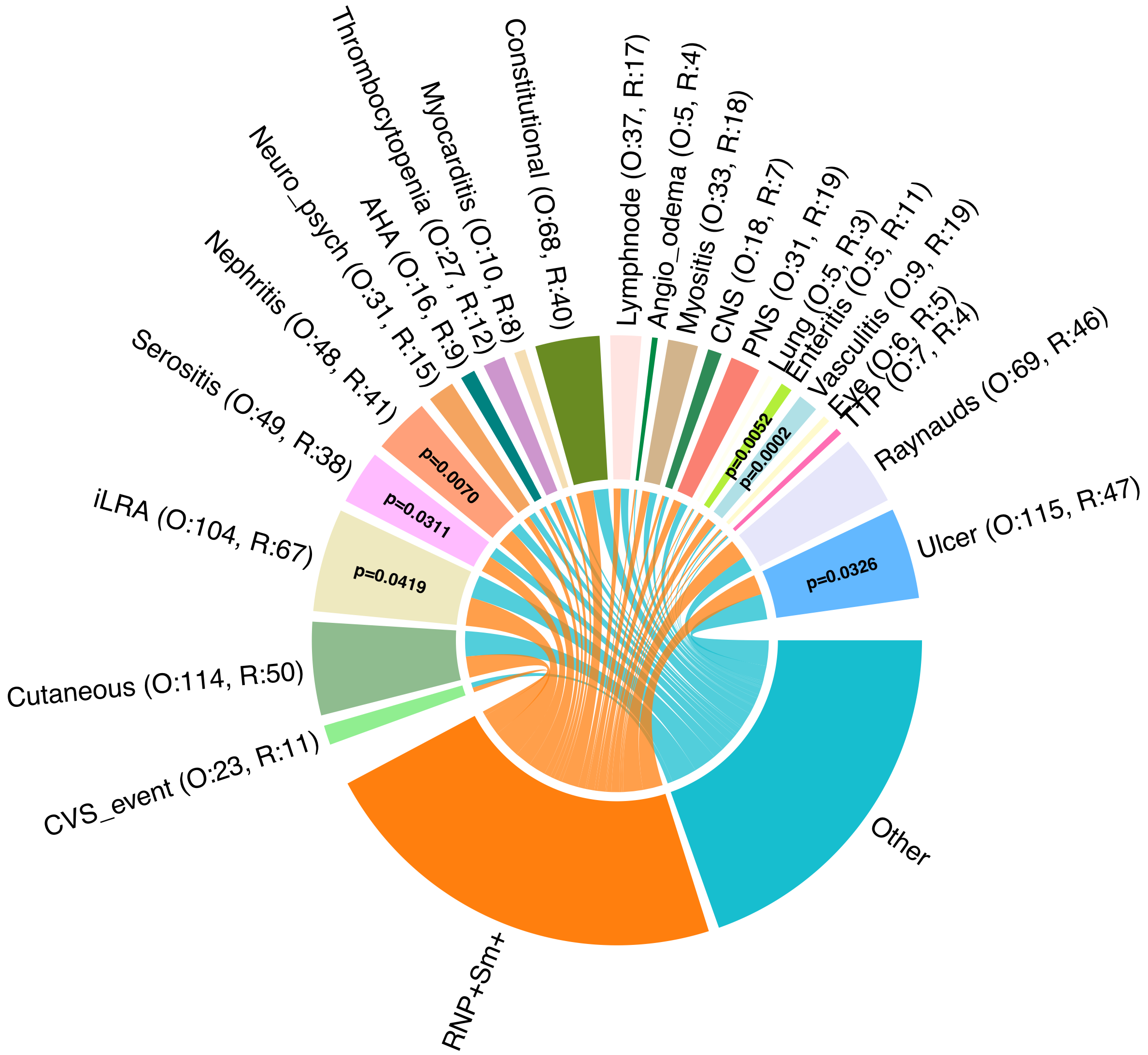


**Supplementary Figure 6: Proportion of patients with various clinical manifestations in lupus patients with anti-RNP and anti-Sm antibodies ($RNP^+Sm^+$) compared with other lupus patients**, using combined data from the BEAT-Lupus and CALIBRATE randomised controlled trials and an observational internal cohort[$]. For each feature, the number of patients is shown (R = $RNP^+Sm^+$, O = Other). p values were calculated using Fisher's exact test and are displayed only when $p < 0.05$.

*[$] Participants were drawn from three cohorts: (i) the BEAT-Lupus clinical trial ($RNP^+Sm^+$, n = 17; Other, n = 35); (ii) CALIBRATE trial ($RNP^+Sm^+$, n = 12; Other, n = 31); and (iii) an internal observational cohort comprising new patients seen at the lupus clinic between June 2019 and June 2024 ($RNP^+Sm^+$, n = 69; Other, n = 122).*

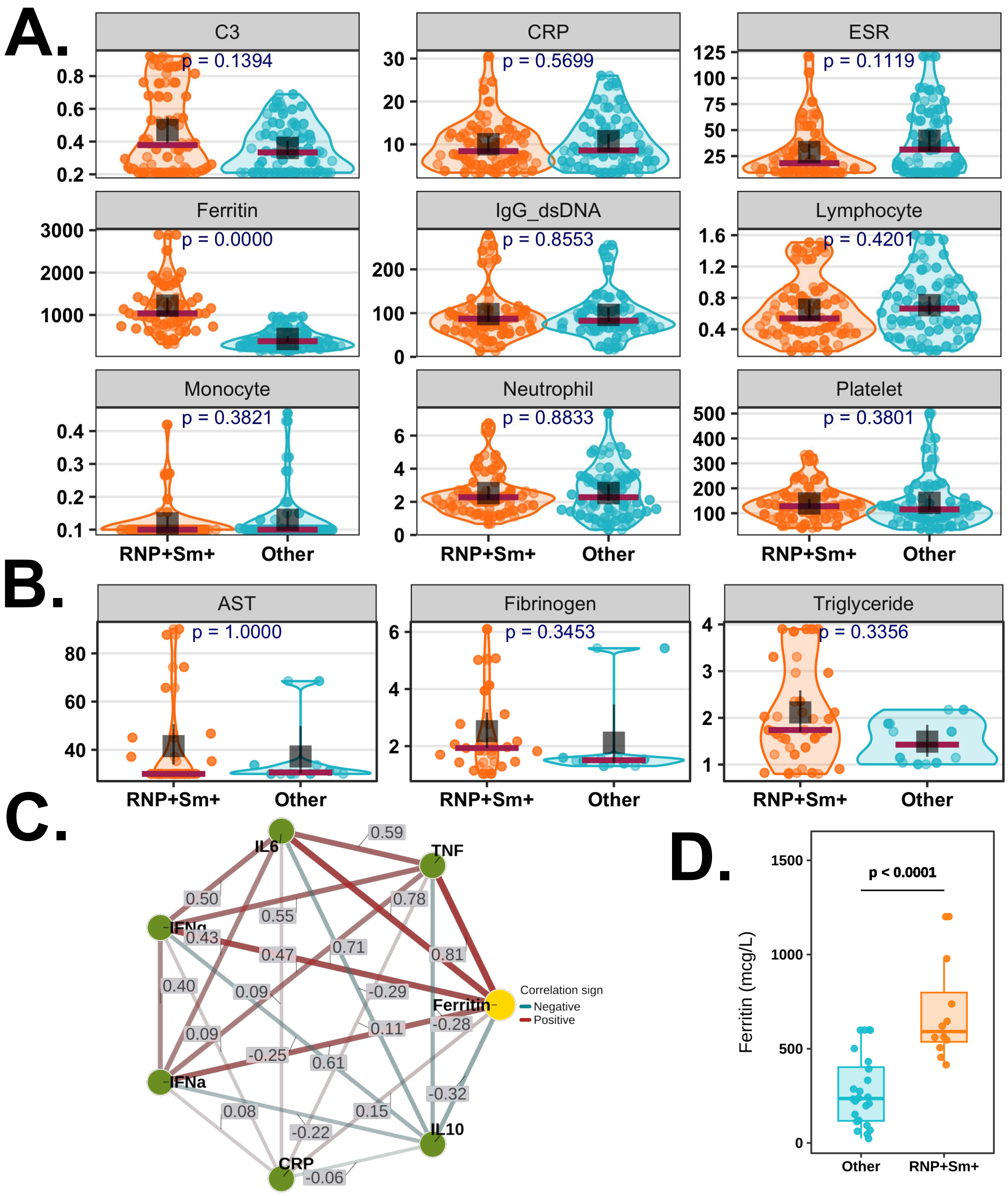


**Supplementary Figure 7 A-D: Conventional laboratory markers and macrophage activation signatures at flare (UCLH and CUH cohorts). A.** Comparison of conventional laboratory markers at the time of flare in the internal UCLH cohort, stratified by endotype: RNP$^+$Sm$^+$ (positive for both anti-RNP and anti-Sm antibodies; $n$ = 42) versus other SLE endotypes ($n$ = 42). **B.** Comparison of additional macrophage activation syndrome (MAS) markers in a subset of flaring patients with available data (RNP$^+$Sm$^+$, $n$ = 21; Other, $n$ = 6). **C.** Correlations (Spearman's) of ferritin with plasma TNF$\alpha$, IL-6, IFN (Interferon)-$\gamma$, and IFN-$\alpha$

levels in the subset of patients with proteomic data (as shown in Figure 1). **D.** Validation of serum ferritin levels in patients with active disease from the CUH cohort ($RNP^{+}Sm^{+}$, $n$ = 12; Other, $n$ = 24). $p$-values were calculated using a linear regression model adjusted for disease activity (SLEDAI), age, and ethnicity.

*C3= serum C3 in g/L, CRP = serum C-reactive protein in mg/L, ESR = Erythrocyte sedimentation rate in mm, Ferritin in mmol/L, IgG_dsDNA = IgG anti-dsDNA antibody in IU/ml, Lymphocyte, monocyte, neutrophil, platelet in x$10^{9}$/L, AST = Aspartate transaminase in IU/L, Fibrinogen in g/L, Triglyceride in mmol/L*

A.

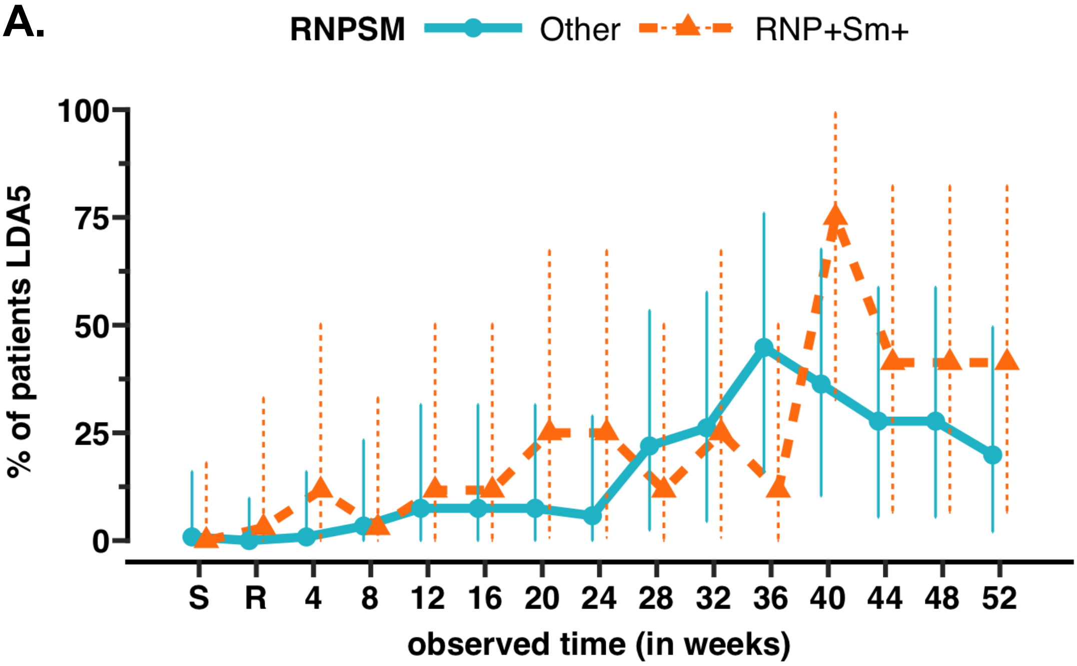


B.

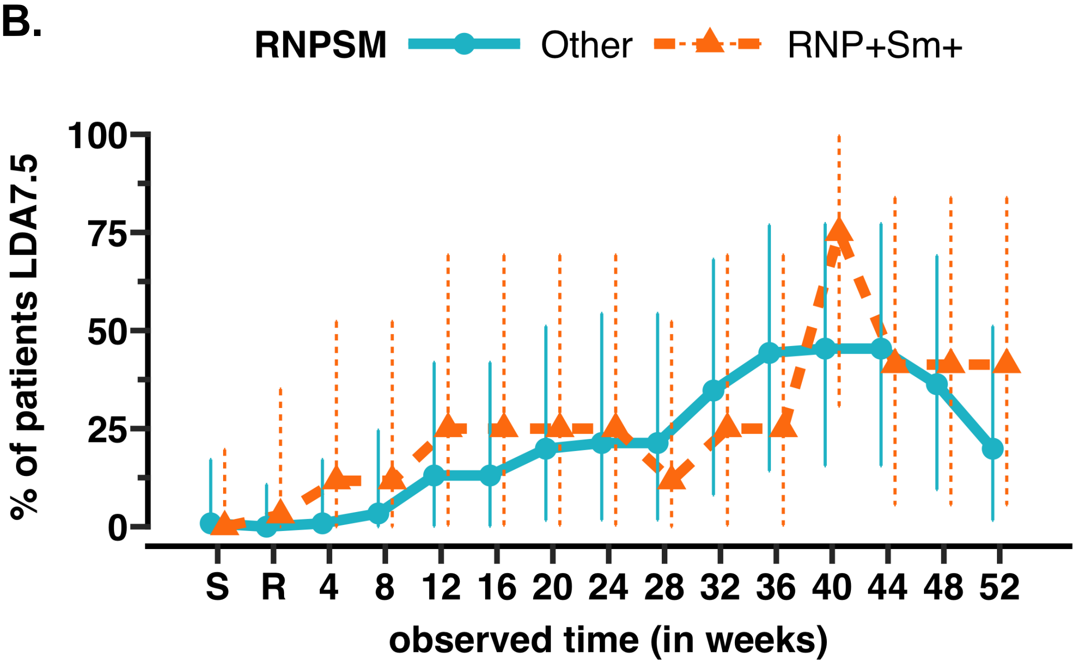


**Supplementary Figure 8A–B: Proportion of patients achieving low disease activity on A. ≤5 mg/day and B. ≤7.5 mg/day prednisolone (LDA5)$^{\$}$, stratified by RNP$^{+}$Sm$^{+}$ endotype (positive for both anti-RNP and anti-Sm antibodies) versus all other patients in the Belimumab arm (n =26) of BEAT-Lupus trial.** (S = screening (6-8 weeks prior randomisation), R = randomisation). $^{\$}$LDA is defined as no BILAG-2004 Grade A or B scores in any organ system at 2 consecutive visits 4 weeks apart (all organ systems scored as C, D, or E), with prednisone dose ≤5 mg/day (LDA5) or ≤7.5 mg/day (LDA7.5) at both visits, stable immunosuppressant and hydroxychloroquine dose for the preceding 12 weeks.

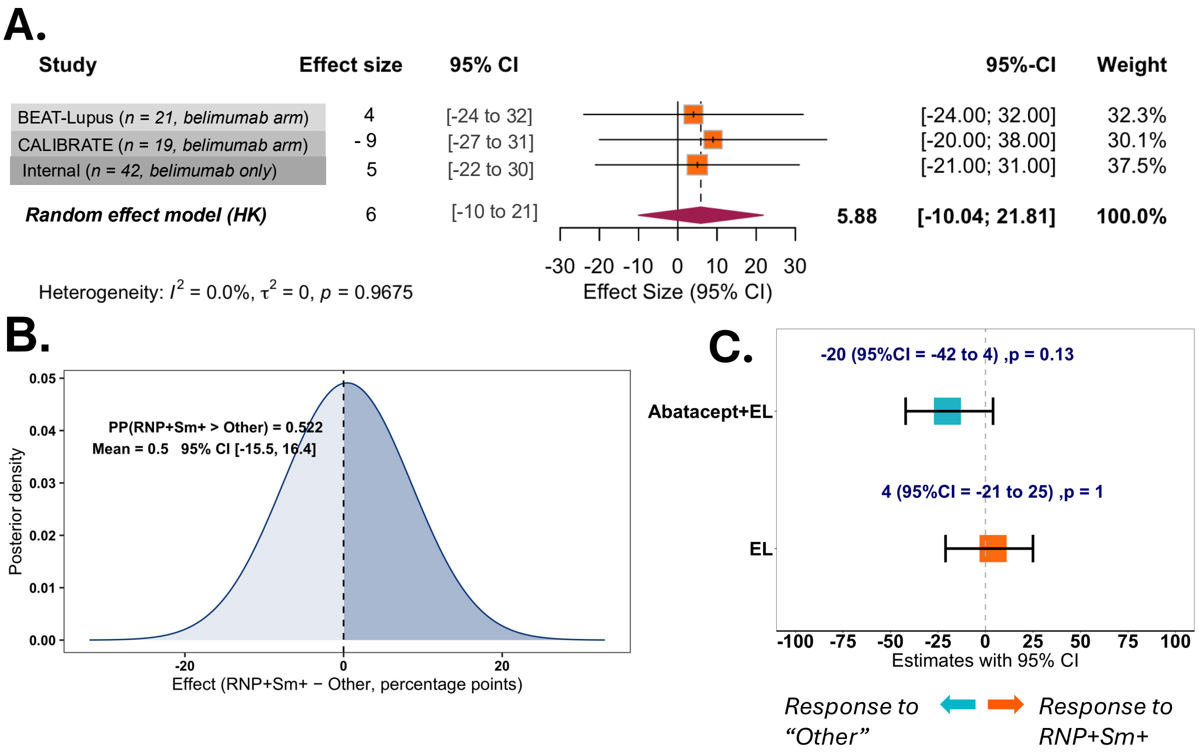


**Supplementary Figure 9 A–C: RNP$^{+}$Sm$^{+}$ (positive for both anti-RNP and anti-Sm antibodies) endotype does not differentiate response to belimumab or cyclophosphamide. A.** Clinical Response Belimumab: The forest plot depicts response rates to belimumab in the BEAT-Lupus and CALIBRATE trials (belimumab was given after rituximab in both trial), and a separate internal observational cohort (treated with belimumab only), evaluated at 52 weeks, 48 weeks, and 24 weeks, respectively. Patients in the observational cohort were treated with belimumab as part of standard NHS England care alongside other therapies. p-values shown here calculated by logistic regression and adjusted for age, ethnicity (Black vs Non-Black) and lupus-nephritis. **B.** Bayesian posterior for the pooled absolute difference in response. Curve shows the posterior density of the overall mean effect (μ) from a Normal–Normal random-effects meta-analysis (flat prior on μ; half-Cauchy(0, 0.5) prior on τ) across the 3 cohorts. Shaded curves are posteriors from a beta–binomial model with Jeffreys prior $\mathrm{Beta}(0.5{,}0.5)$ fitted to pooled responder/non-responder counts from the three studies. **C.** Treatment and endotype interaction effects across 3 RCTs. Forest plot illustrating stratum-specific odds ratios (ORs) for clinical response to active versus control regimens, stratified by endotype: RNP$^{+}$Sm$^{+}$ vs Other. ORs and 95% Wald confidence intervals were estimated by penalised logistic regression with a treatment × endotype interaction (Bayesian weakly-informative prior). p-values shown here calculated by logistic regression and adjusted for age, ethnicity (Black vs Non-Black) and lupus-nephritis.

*Trial/regimen contrasts: BEAT (RB: rituximab+belimumab vs. R: rituximab alone, week 52), CALIBRATE (RCB: rituximab–cyclophosphamide+belimumab vs. RC: rituximab–cyclophosphamide, week 48), and ACCESS (Euro-Lupus AC: cyclophosphamide±abatacept, vs cyclophosphamide only: week 24).*

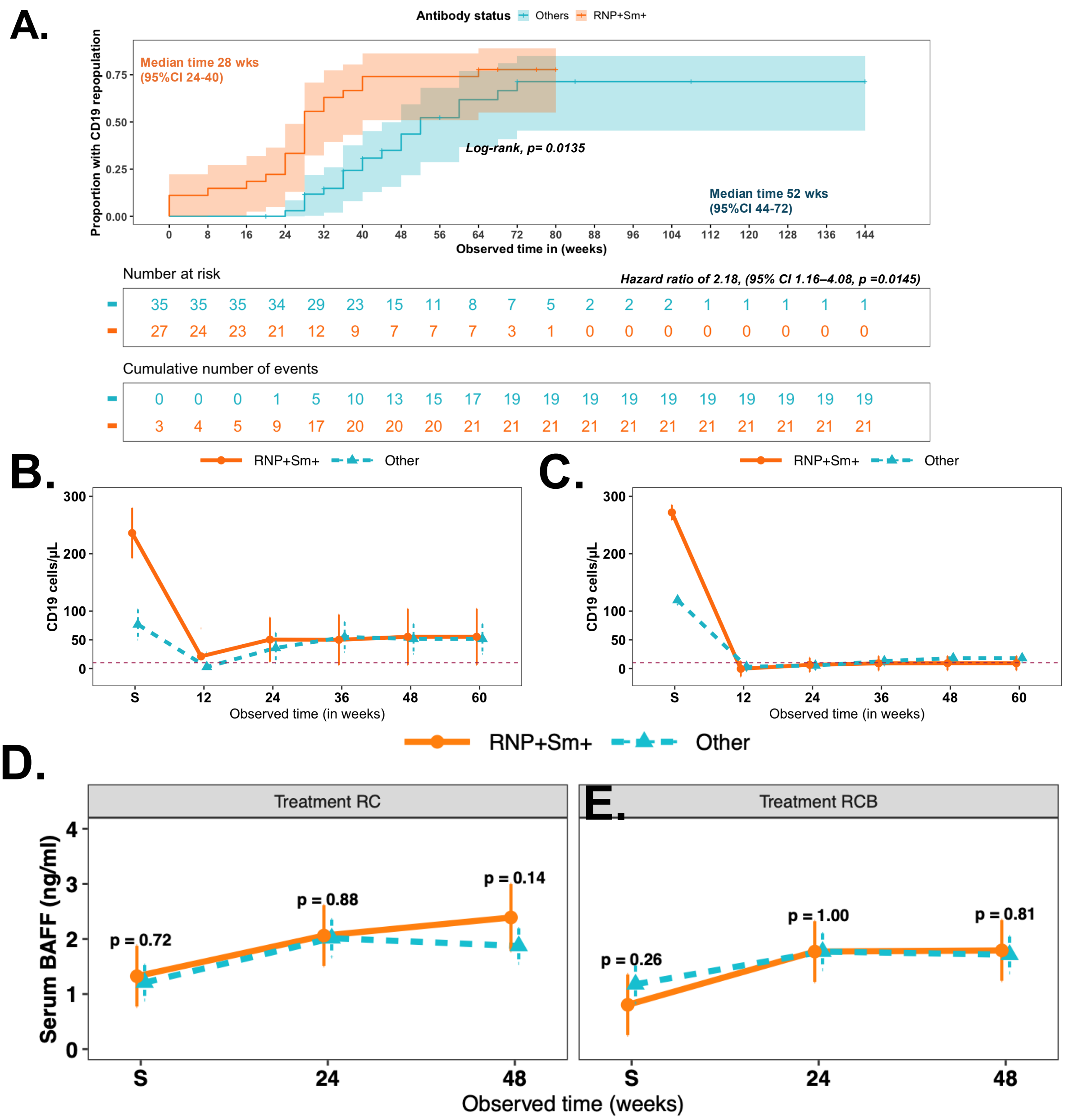


**Supplementary Figure 10 A–E: Changes in CD19$^+$ B-cell counts and serum BAFF (B-cell activating factor) stratified by endotype (RNP$^+$Sm$^+$: positive for both anti-RNP and anti-Sm) versus all other patients. A.** UCLH internal observational cohort: Kaplan–Meier curves for time to CD19$^+$ B-cell repopulation; Cox regression adjusted for age, ethnicity, and medication dose. **B–E.** CALIBRATE trial: Changes in CD19$^+$ B cells in **B.** rituximab–cyclophosphamide (RC) arm and **C.** rituximab–cyclophosphamide–belimumab (RCB) arm, from screening (S) to week 60. Horizontal dashed line marks 10 cells/μL (B-cell depletion threshold). Serum BAFF changes in **D.** RC arm and **E.** RCB arm, from screening (S) to week 48. Lines show geometric means (95% CIs) by group over time, estimated from a mixed-effects model with a log link (random intercept for participant; fixed effects for group, time, and their interaction). Estimated marginal means were back-transformed to the original scale. p-values were calculated using Tukey-adjusted pairwise contrasts between groups at each visit, with adjustment for age, ethnicity, baseline medication dose, and lupus nephritis status.

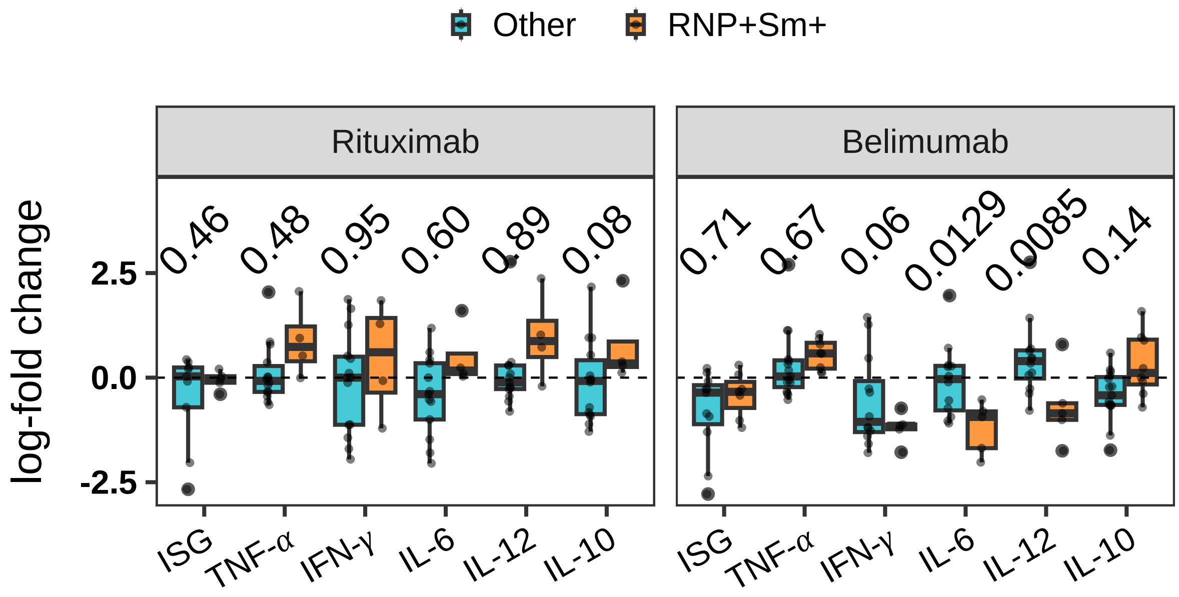


**Supplementary Figure 11: Changes in interferon-stimulated gene (ISG) expression and baseline-elevated cytokines in the Beat-Lupus trial at 12 months from baseline, stratified by autoantibody subgroup and treatment arm.** Panels show the log fold-change from baseline to month 12 for ISGs and cytokines that were elevated at baseline (Supplementary Figure 1), analysed separately for the Rituximab/Placebo arm and the Belimumab arm. Patients were stratified into RNP$^{+}$Sm$^{+}$ (Rituximab: n = 5; Belimumab: n = 9) and Other autoantibody groups (Rituximab: n = 14; Belimumab: n = 14). p-values were calculated using ANCOVA on log-transformed Week 52 values, adjusted for baseline levels, age, ethnicity, and baseline SLEDAI, comparing the RNP$^{+}$Sm$^{+}$ subgroup to the Other subgroup. Multiple comparisons were corrected using Tukey's post hoc test.

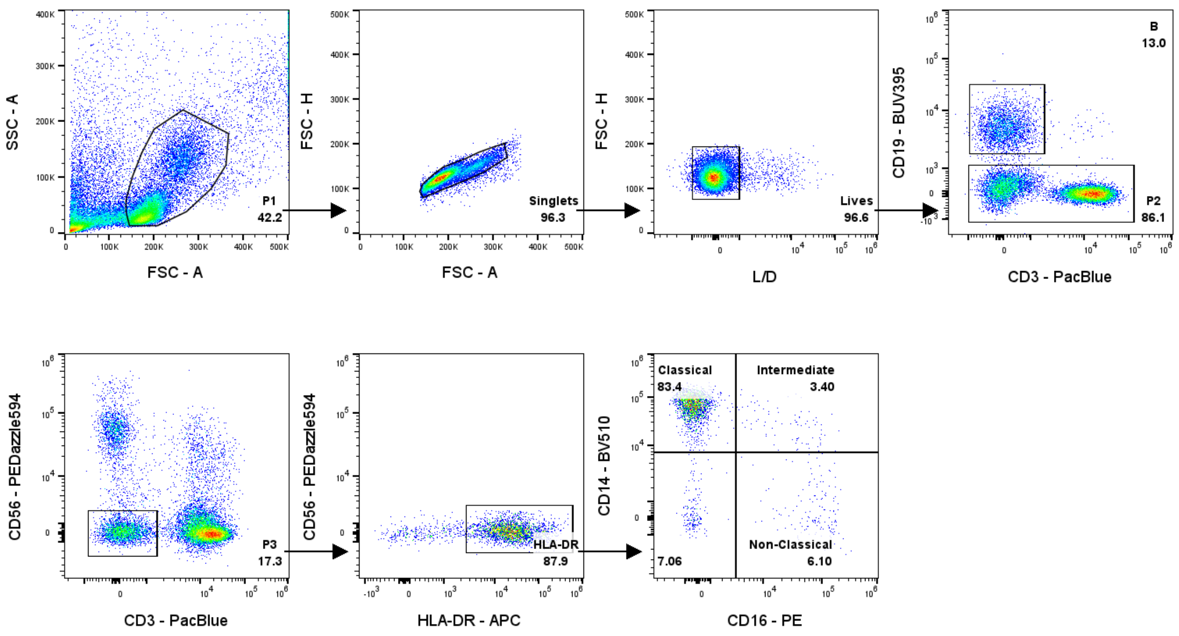


**Supplementary Figure 12:** Gating strategies to define intermediate, non-classical and classical monocytes.

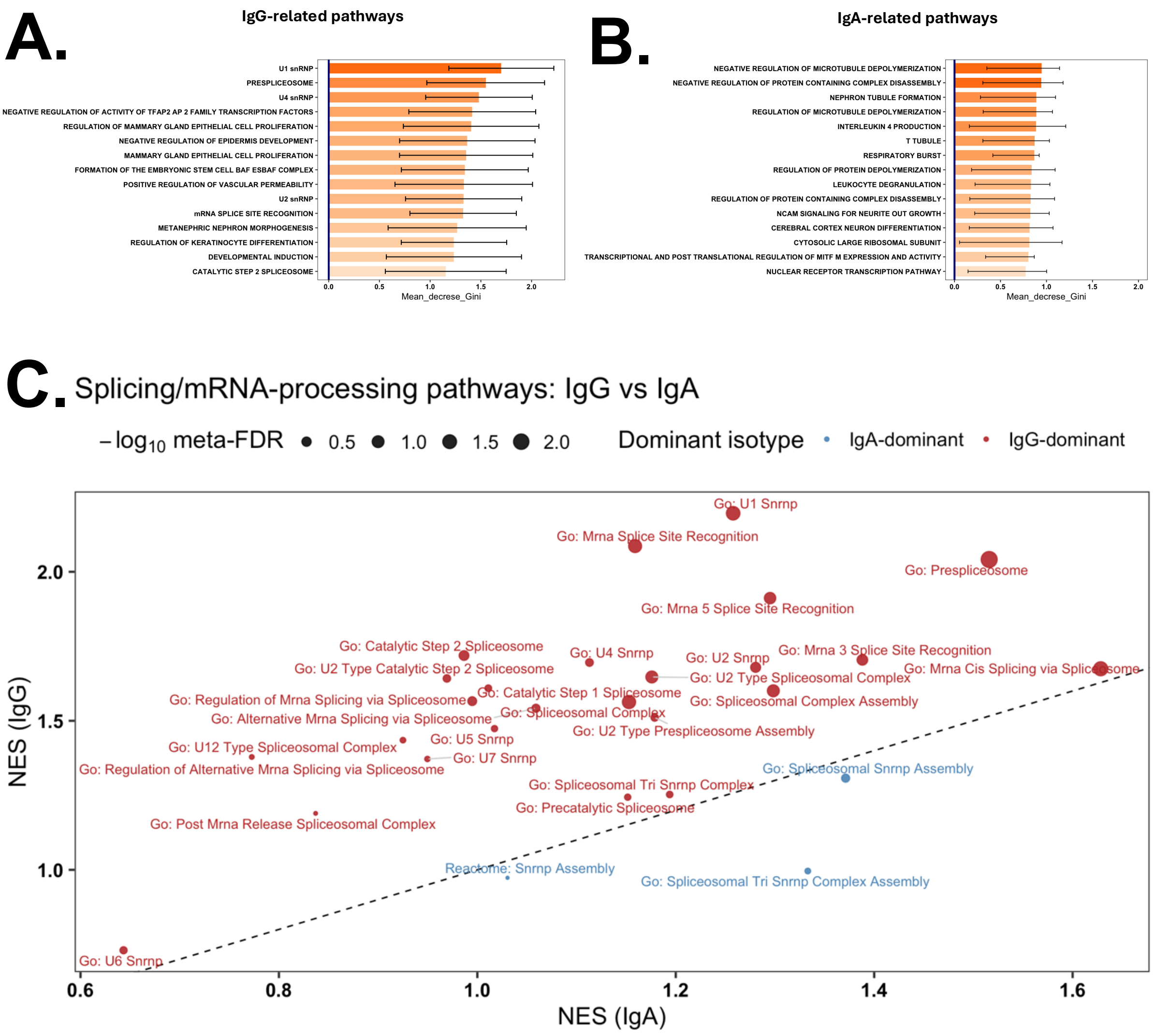


**Supplementary Figure 13A–C: IgG (but not IgA)-associated autoreactivity predominates in spliceosome-related modules in RNP⁺Sm⁺ patients (BEAT-Lupus cohort).** Bar plots display **B.** IgG and **C.** IgA-associated autoreactivity scores of the top fifteen significantly enriched functional sets identified by pre-ranked gene set enrichment (cameraPR) analysis using MSigDB annotations (GO:CC, GO:BP, Reactome). Bars show the mean difference in module scores between $RNP^+Sm^+$ (n=9) and other patients (n=11) ($\Delta = RNP^+Sm^+ - \text{Other}$), with 95% confidence intervals, adjusted using corresponding healthy controls from each batch. **C.** Scatter plot displaying gene set enrichment analysis (GSEA) normalised enrichment scores (NES) for $RNP^+Sm^+$ versus Other across pathways annotated to splicing/mRNA processing (GO:BP, GO:CC, Reactome). The x-axis shows NES based on IgA-associated ranks; the y-axis shows NES from IgG-associated ranks. The dashed line (y = x) indicates equal enrichment in both isotypes. Each point represents a single pathway, with point size denoting $-\log_{10}$(meta-FDR) from a Z-score meta-analysis across isotypes.

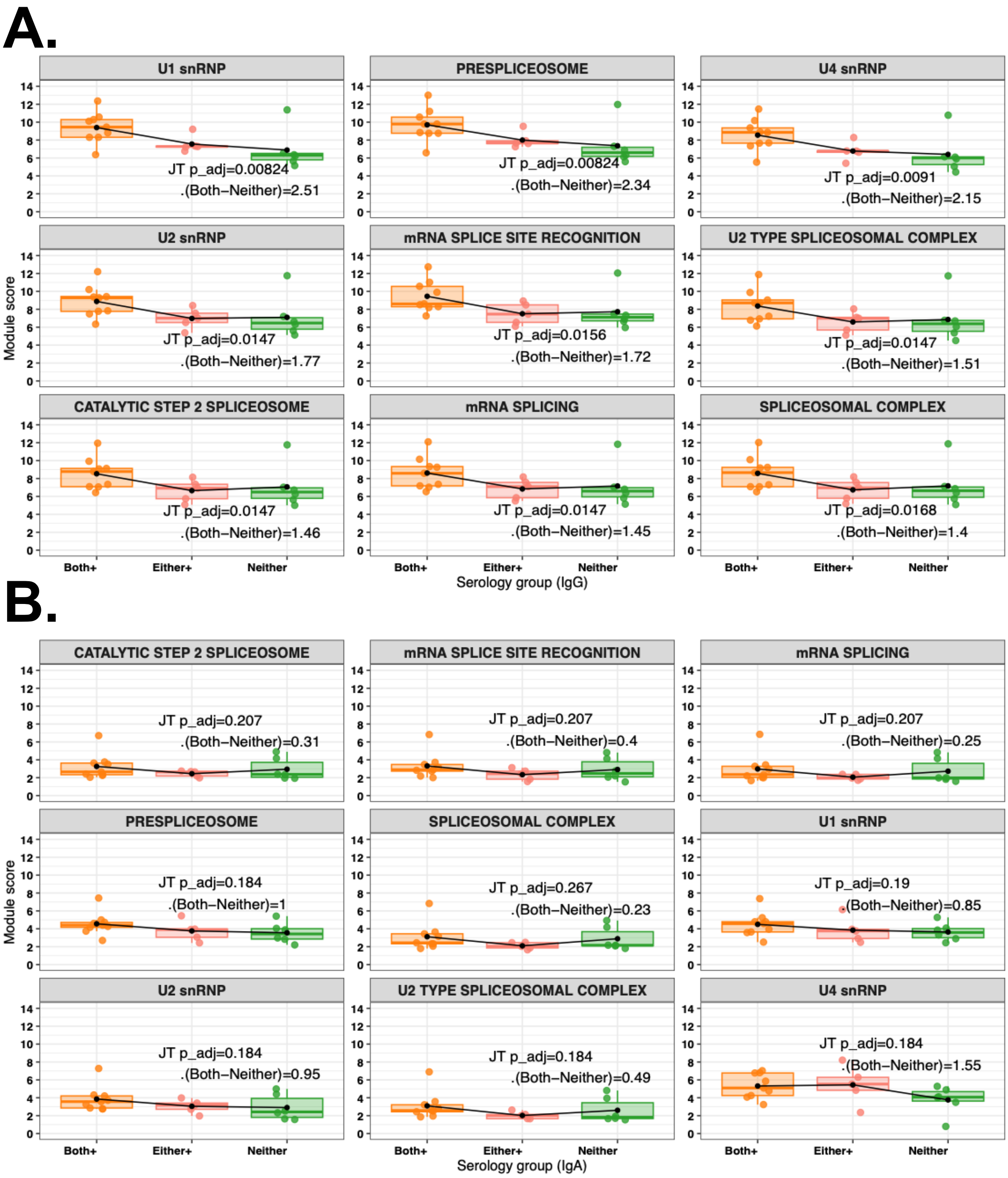


**Supplementary Figure 14A–B: Jonckheere–Terpstra (JT) trend analysis of A. IgG-and B. IgA- associated spliceosome modules across ordered serology groups**: RNP$^{+}$Sm$^{+}$ (Both+), n=9; RNP- or Sm-single positive (Either+), n=5; and seronegative (Neither), n=6. Box and jitter plots display individual patient module scores, with lines indicating group means. Adjusted p-values for JT tests and effect sizes (Δ Both+ – Neither) are annotated.

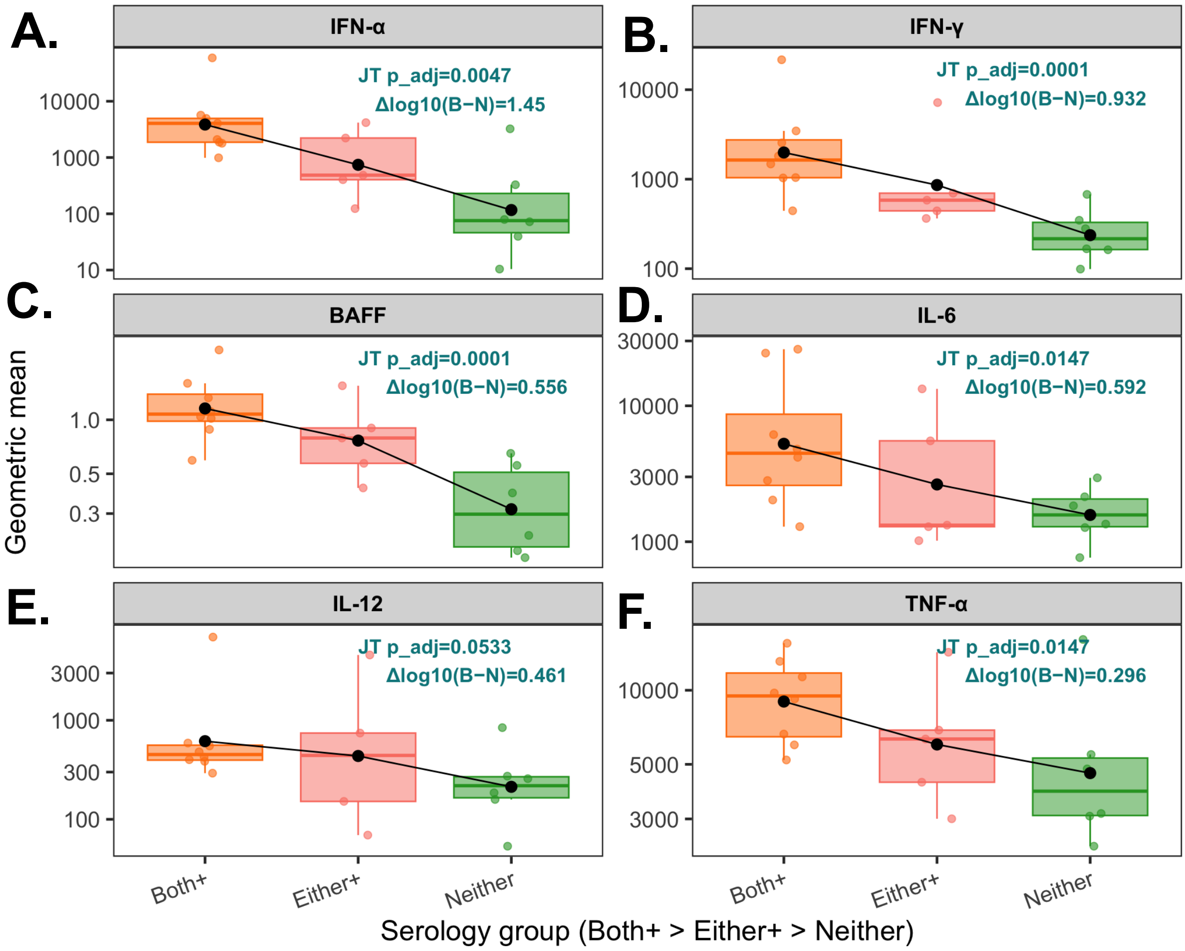


**Supplementary Figure 15 A–F: Jonckheere–Terpstra (JT) trend analysis of 6 key cytokines of those patients with spliceosome modules available across ordered serology groups:** RNP$^{+}$Sm$^{+}$ (Both+), n=9; RNP- or Sm-single positive (Either+), n=5; and seronegative (Neither), n=6. Box and jitter plots display individual patient module scores, with lines indicating group means. Adjusted p-values for JT tests and effect sizes (Δ Both+ – Neither) are annotated.

2. Atisha-Fregoso Y, Malkiel S, Harris KM, Byron M, Ding L, Kanaparthi S, et al. Phase II Randomized Trial of Rituximab Plus Cyclophosphamide Followed by

Belimumab for the Treatment of Lupus Nephritis. Arthritis Rheumatol. 2021;73(1):121-31.
3. Group TAT. Treatment of Lupus Nephritis With Abatacept: The Abatacept and Cyclophosphamide Combination Efficacy and Safety Study. Arthritis & Rheumatology. 2014;66(11):3096-104.
